# The first near-complete genome assembly of pig: enabling more accurate genetic research

**DOI:** 10.1101/2024.10.13.617951

**Authors:** Caiyun Cao, Jian Miao, Qinqin Xie, Jiabao Sun, Hong Cheng, Zhenyang Zhang, Fen Wu, Shuang Liu, Xiaowei Ye, Zhe Zhang, Qishan Wang, Yuchun Pan, Zhen Wang

**Affiliations:** College of Animal Sciences, Zhejiang University, Hangzhou, Zhejiang 310058, China; Hainan Institute of Zhejiang University, Building 11, Yongyou Industrial Park, Yazhou Bay Science and Technology City, Yazhou District, Sanya, 572025, Hainan, China

**Keywords:** Pig genome assembly, HiFi and ONT sequencing, gapless reference genome, telomere-to-telomere

## Abstract

Pigs are crucial sources of meat and protein, valuable animal models, and potential donors for xenotransplantation. However, the existing reference genome for pigs is incomplete, with thousands of segments and missing centromeres and telomeres, which limits our understanding of the important traits in these genomic regions. To address this issue, we present a near complete genome assembly for the Jinhua pig (JH-T2T), constructed using PacBio HiFi and ONT long reads. This assembly includes all 18 autosomes and the X and Y sex chromosomes, with only six gaps. It features annotations of 46.90% repetitive sequences, 35 telomeres, 17 centromeres, and 23,924 high-confident genes. Compared to the Sscrofa11.1, JH-T2T closes nearly all gaps, extends sequences by 177 Mb, predicts more intact telomeres and centromeres, and gains 799 more genes and loses 114 genes. Moreover, it enhances the mapping rate for both Western and Chinese local pigs, outperforming Sscrofa11.1 as a reference genome. Additionally, this comprehensive genome assembly will facilitate large-scale variant detection and enable the exploration of genes associated with pig domestication, such as *GPAM*, *CYP2C18*, *LY9*, *ITLN2*, and *CHIA*. Our findings represent a significant advancement in pig genomics, providing a robust resource that enhances genetic research, breeding programs, and biomedical applications.

## Introduction

Pig (*Sus scrofa*) is not only economically important due to its role as a food source but also serves as a medical model and xenotransplantation donor because of its anatomical and physiological similarities with humans [1,2]. Understanding the genome and gene content of candidate species, including pigs, is crucial for selecting the best animal model species for pharmacological or toxicological studies. High-quality, fully annotated genome sequences are essential for gene editing, producing improved animal models for research, or providing cells and tissues for xenotransplantation, as well as enhancing productivity [3,4].

Despite the availability of several high-quality pig reference genomes, including those of the European Duroc [5], Ninxiang [6], Meishan [7], and Jinhua [8,9] pig genomes, these assemblies remain incomplete in genomic regions of repetitive sequences, centromeres, and telomeres [5–9]. A gap-free genome is the ultimate goal of genome assembly, crucial for improving the accuracy of read mapping and variant calling for individuals sequenced with short and long reads [10], and offers new opportunities for identifying unique genes and structural variations (SVs) [11,12]. However, to date, a gapless pig reference genome has not yet been reported.

Advancements in new sequencing technologies and computational algorithms have ushered in the era of telomere-to-telomere (T2T) assemblies [13]. Specifically, third-generation sequencing technologies, which generate long reads enabling whole-genome assembly, have improved both experimental methods and algorithms. For example, Pacific Biosciences (PacBio) methods can generate ∼10 Kb long HiFi reads with 99% accuracy, while Oxford Nanopore Technologies (ONT) recently developed an ultra-long read method producing reads with an average length of ∼50 Kb, extending up to ∼100 Kb, with the longest reads reaching hundreds of Kb [14–16]. HiFi reads can assist in assembling complex genomic regions[17], while the ONT ultra-long reads can help assemble genomic regions with tandem duplications[18]. The application of third-generation sequencing and assembly technologies to high-fidelity long reads has contributed to the creation of gap-free genome assemblies across hundreds of species [19].

Therefore, in response to the gap of a T2T-level pig genome, we assembled a gap-free T2T genome of the Jinhua pig-one of China’s four renowned indigenous breeds, famous for its superior meat quality and high-quality Jinhua-ham [20] using PacBio HiFi and ONT long reads. This T2T genome assembly marks a significant advancement in pig genomics. It offers enhanced resources for research in pig genetics, genomics, and biomedical applications. This assembly overcomes the limitations of previous incomplete assemblies, serving as a robust platform for various downstream comparative genomic analyses and providing new insights into the complex traits of pigs.

## Results

### T2T assembly of JH pig genome

We generated a total of 51.10× sequence coverage of raw PacBio HiFi data (135.95 Gb, read N50 18.32 Kb), 136.65× sequence coverage of ultralong ONT data (363.55 Gb, read N50 52.17 Kb), 94× sequence coverage of Hi-C data, and 50× sequence coverage of WGS data for assembling the JH pig genome (**Figure S2A-D** and **Supplemental Table S1**). Using the HiFi reads, we assembled the initial PacBio HiFi assembly, which had a total length of 2.72 Gb and 187 contigs (contig N50 84.83 Mb, **Figure S1E**). The initial ONT assembly had a total length of 2.28 Gb and 93 contigs (contig N50 64.34 Mb, **Figure 1A, Figure S2F,** and **Supplemental Table S1**). A second ONT assembly using only the longest ONT reads (87.23G, read N50 100 Kb) had a total length of 2.31 Gb and 112 contigs (contig N50 72.01 Mb, **Figure S2G**), which were used to fill the gaps at the later polish steps. Since the PacBio HiFi assembly showed a higher quality and contiguity compared to the ONT assembly, we selected it for the bone genome assembly. We used Hi-C data to order and orient these PacBio HiFi contigs, resulting in 20 chromosomes (with six gaps, scaffold N50 142.74 Mb) representing chromosomes 1-18, X, Y, and 66 unplaced contigs containing an additional 62.32 Mb (**Figure 1A-B, Figure S2E,** and **Supplementary Table S3)**. The PacBio HiFi assembly was further iteratively polished by PacBio HiFi reads, ONT reads, Hi-C data (for mapping error correction), and the second ONT assembly, resulting in a near-T2T assembly with a total length of 2.68 Gb (2.61 Gb mounted on the chromosome, mounting rate of 97.67%, contig N50 142.75 Mb) and only six gaps remaining in chromosomes 2, 3, 8, and 10 (**Figure S2E** and **Supplementary Table S3**).

**Figure 1.**
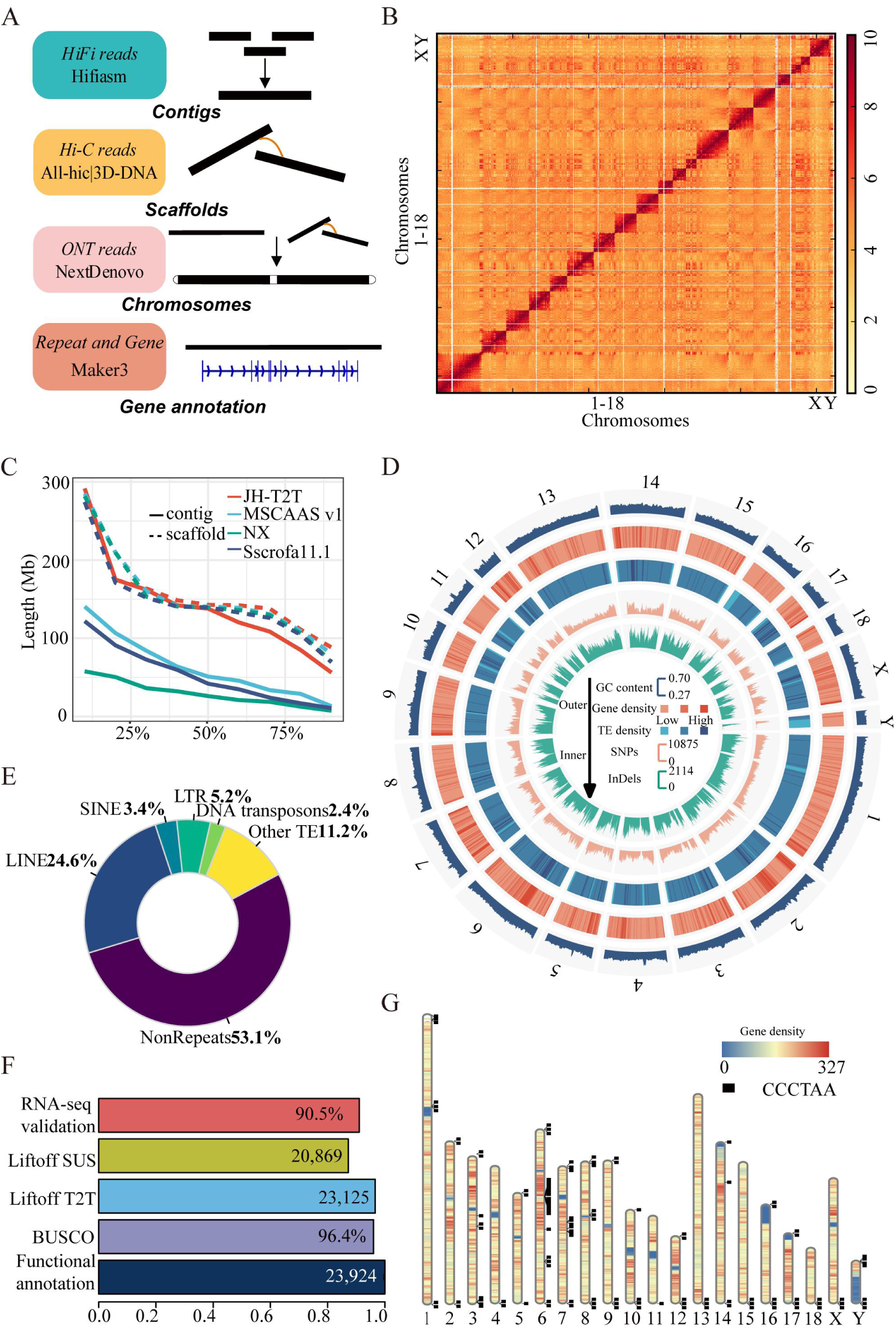
Summary of JH-T2T pig genome assembly. (A) Schematic diagram illustrating the pipeline for genome assembly and annotation. (B) Hi-C chromatin interactions of the assembled JH-T2T genome. (C) Comparison of the contiguity between released assemblies and JH-T2T assembly. (D) Landscape of the assembled JH-T2T genome, showing chromosomes, GC contents, gene, repeat and TE density, SNPs, and InDels in different tracks from outer to inner. (E) Composition ratio of repeat elements in JH-T2T. (F) Gene annotations of JH-T2T. (G) Genome-wide telomere portrait of JH-T2T. The black boxes indicate chromosomal loci of the tandemly repeated telomeric motif in the primary assembly. The heatmap shows the chromosome-wide repeat density in non-overlapping 1 Mb windows.

### Quality assessment of the final JH-T2T assembly

We conducted a comprehensive assessment of the JH-T2T assembly’s quality and completeness in multiple ways. First, the estimated chromosome size was determined to be 2.69 Gb, with a heterozygosity rate of 0.38%, consistent with the Sscrofa11.1 genome size (**Figure S4A** and **Supplementary Table S3**). Second, 16 of the 20 chromosomes were each represented by a single contig (**Figure 1D** and **Supplementary Table S3**), indicating superior sequence integrity compared to the current pig reference genome, Sscrofa11.1 (1,117 contigs), and other published pig genomes (**Figure 1C**, **Table 1**, and **Supplementary Table S4)**. Third, the JH-T2T assembly showed high overall base accuracy, estimated at 99.997% (an average QV score of 55) using mapped K-mers from trio Illumina PCR-free reads data. QV scores ranged from 48 to 62 for each chromosome, with five chromosomes (chr4, chr9, chr11, chr15 and chr16) having high QV scores greater than 60 (**Figure S4C-D** and **Supplementary Table S3**). Forth, compared to the other three genomes, BUSCO analysis revealed that the JH-T2T assembly exhibited the highest percentage of completeness, with approximately 96.4% of the core conserved mammalian genes being fully represented (**Figure 1F, Figure S4B** and **Supplementary Table S5**). This indicates a near-complete genome assembly. Fifth, the chromosomal interaction maps generated using Hi-C data provided further evidence of the accuracy and reliability of the JH-T2T assembly. Hi-C data revealed that all chromosomes displayed clear intra-chromosomal diagonal signals, with no significant inter-chromosomal signals, confirming the correct order and orientation of all pseudomolecules (**Figure 1B** and **Supplementary Table S2**). Sixth, the remapping rates for HiFi reads, ONT reads, and Illumina short reads on JH-T2T assembly were impressively high at 99.90%, 99.99%, and 99.99%, respectively.

**Table 1.** Summary information of JH-T2T, Sscrofa11.1, Ningxiang and MSCAAS v1 assemblies.

| Terms | JH-T2T | Sscrofa11.1[5] | Ningxiang[6] | MSCAAS v1[7] |
| --- | --- | --- | --- | --- |
| Contig N50 (Mb) | 100.5 | 48.2 | 26.1 | 48.1 |
| Contig number | 26 | 1,117 | 305 | 152 |
| Scaffold N50 (Mb) | 142.7 | 88.2 | 139.0 | 139.0 |
| Scaffold number | 20 | 20 | 19 | 19 |
| Gaps | 6 | 103 | 286 | 133 |
| Assembly size (Gb) | 2.61 | 2.50 | 2.44 | 2.50 |
| Average length of CDS (bp) | 1,593 | 1,668 | 1,601 | 1,379 |
| Protein-coding genes | 23,924 | 20,661 | 20,914 | 22,855 |

For alignment-based comparison with other reported genomes, we firstly utilized WGS data from 30 individuals (depth ranging from 10.00 to 27.14×, **Supplemental Table S6**), which were mapped to the Sscrofa11.1 [5], MS [7], NX [6], and JH-T2T genomes. The JH-T2T showed significantly higher mapping rates ranging (98.65% to 99.87%, **Figure 2A**), properly mapped rate (92.16% to 98.51%, **Figure 2B**) and lower Base error rates (0.64% to 1.62%, **Figure 2C**) compared to other genomes. The average mapping rate for Asian pigs was 99.53% on JH-T2T versus 97.98% on Sscrofa11.1, and for European pigs, 99.48% versus 98.44% (**Figure 2A**). The average properly mapped rate for Asian pigs was 97.84% on JH-T2T versus 94.68% on Sscrofa11.1, and for European pigs, 94.78% versus 93.04% (**Figure 2B**). Next, mapping 111 RNA-seq data (**Supplemental Table S7**) from Asian (n=61) and European (n=50) pig breeds showed that JH-T2T was more suitable for analyzing RNA-seq data from Asian pig breeds, with higher mapping rates (88.70%) compared to Sscrofa11.1 (87.67%, **Figure 2D**). The average mapping rate of European pigs on JH-T2T and Sscrofa11.1 was similar (89.45% versus 89.58%, **Figure 2D**). The above results suggest that JH-T2T will be advantageous for both DNA and RNA sequencing data mapping analysis.

**Figure 2.**
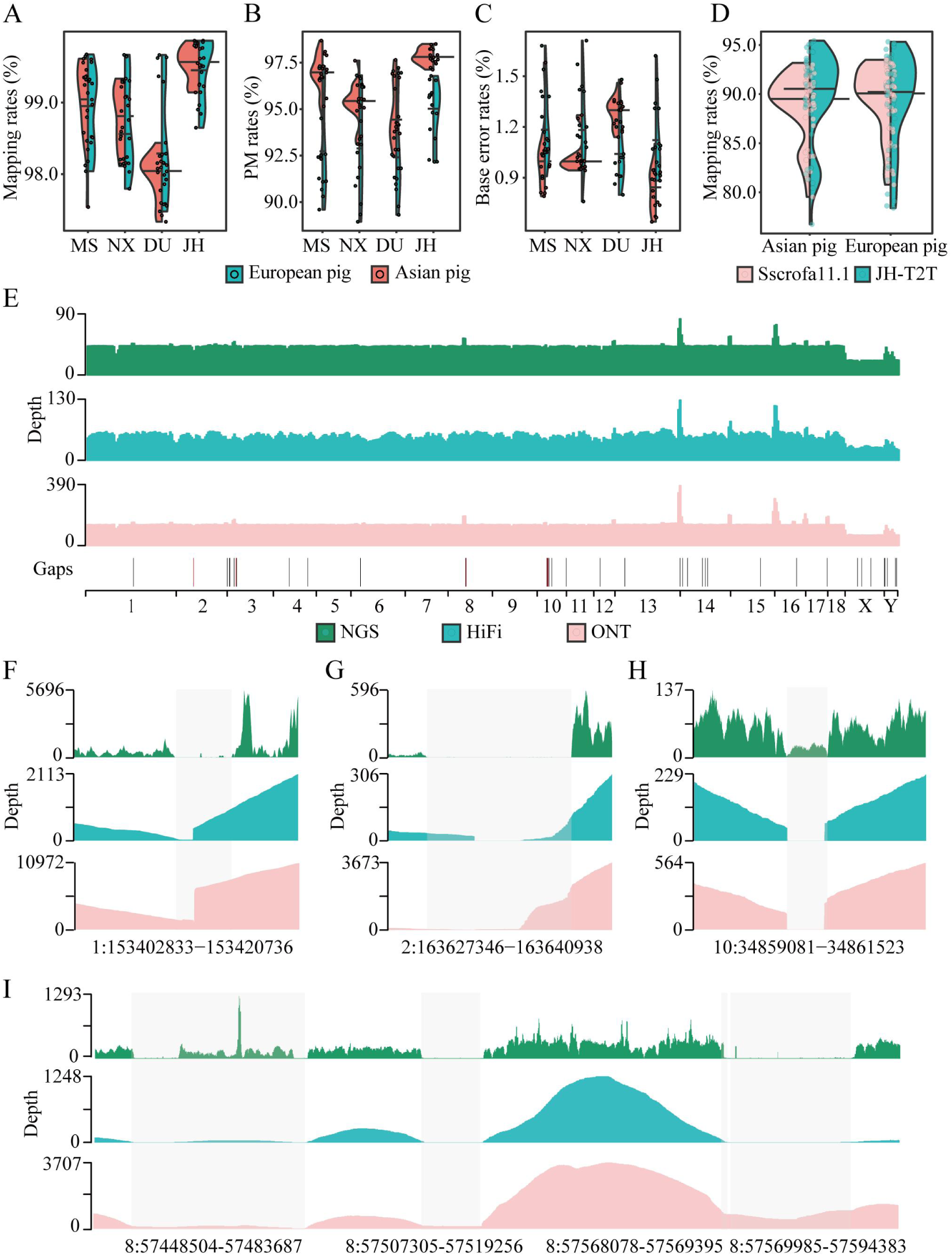
Sequencing coverage, mapping rate and filling gaps in JH-T2T assembly. (A-C) Comparison of DNA sequencing read mapping rates when whole genome resequencing reads from Asian (Left) and European (Right) pig mapped to MSCAAS v1, Ningxiang (NX), Sscrofa11.1 and JH-T2T genome assemblies, respectively. (D) Comparison of RNA sequencing read mapping rates for Asian (Left) and European (Right) mapped to the Duroc (Sscrofa11.1) and the JH-T2T genome assembly, respectively. (E) Whole-genome sequence coverage of mapped WGS, HiFi and ONT reads. Gaps distribution across chromosomes in JH-T2T. Black indicates filled gaps, while red indicates unfilled gaps. (F-I) Whole-genome sequence coverage of mapped WGS, HiFi and ONT reads specifically in gap regions.

Additionally, 49 out of 63 gaps in the historical genome version were successfully closed and 8 out of 63 gaps were corrected in our final JH assembly, with filled gaps ranged from 81 to 35,183 bp, totaling around 268 Kb (**Figure S3, S5** and **Supplemental Table S8**). We remapped ONT and HiFi reads to the post-gap filling genome to confirm the reliability for each filled gap. Most filled gaps were identifiable through ONT or HiFi alignments, and assembly errors in low-coverage regions (LCRs) were corrected via ONT alignments (**Figure 2E** and **Figure S6**). Specifically, the largest two gaps (36 and 24 Kb) on chromosome 8 were successfully confirmed by coverage with multiple ONT or HiFi reads (**Figure 2I** and **Figure S5-6**). The gaps on chromosomes 1, 2 and 10 were also successfully filled, evidenced by fully coverage with both ONT and HiFi reads (**Figure 2F-H** and **Supplementary Table S11**). These findings confirmed the accuracy and reliability of our JH-T2T assembly.

Overall, our assembly quality metrics indicate a gapless, fully phased, and near-T2T assembly of the pig genome. To the best of our knowledge, this assembly is the first T2T and the most complete pig genome assembly published.

### Genome annotation

The JH-T2T genome assembly provides a gapless T2T sequence for all 20 chromosomes, marking significant progress over previous incomplete pig genome assemblies [5–7]. About 46.90% of the JH-T2T genome consists of repetitive sequences elements: 24.63% LINEs (long interspersed nuclear elements), 3.40% SINEs (short interspersed nuclear elements), 5.23% LTR (long terminal repeat), 2.44% DNA transposons, 1.13% simple repeats, and 5.21% satellites (**Supplemental Table 9** and **Figure 1E**).

Using the six-base telomere repeats (TTAGGG/CCCTAA) as a query, 35 out of the anticipated 40 telomeres were identified, with a single telomere detected on chromosomes 4, 11, 15, 18, and 19. The average telomere length is 17.95 Kb, with approximately 2,039 repeat copies per telomere. The longest telomere spanned 19.99 Kb (**Figure 1G** and **Supplementary Table S3**). Putative centromeres were identified in expected locations on chromosomes 1–12, 14, and 16-17 (**Figure S7** and **Supplementary Table S3**).

Gene annotation of the masked JH-T2T genome was performed using MAKER [37] with evidence from protein homologies and RNA-seq data. A total of 23,924 high-confidence protein-coding genes were predicted (**Figure 1F** and **Supplementary Table S13**), which includes 799 newly anchored genes (**Supplementary Table S17**). To validate these predictions, RNA-seq data from 111 samples showed that 20,110 (90.50%) of the high-confidence genes were expressed in at least one sample (**Figure 1F**). Gene and repeat distribution across chromosomes follow the typical pattern observed in vertebrate genomes, with higher gene concentrations in GC-rich regions and decreased gene density in repeat-rich distal regions (**Figure 1D**). Also, the density of genes at telomeres is lower (**Figure 1G**).

### Global comparison between the Sscrofa11.1 and JH-T2T genome

The JH-T2T genome assembly showcases greater completeness and accuracy compared to the Sscrofa11.1 assembly. First, the JH-T2T assembly added approximately 171 Mb (6.8%) to the Sscrofa11.1 assembly. Second, the completeness measured by BUSCOs, the JH-T2T achieved 96.4% of 9,226 BUSCOs, surpassing Sscrofa11.1’s 94.1% (**Figure S4B** and **Supplementary Table S5**). Third, the JH-T2T assembly identified 35 telomeres (out of an expected 40, **Supplementary Table S3**), whereas Sscrofa11.1 captured telomere only at the proximal ends of Sscrofa11.1 chromosome assemblies of SSC2, SSC3, SSC6, SSC8, SSC9, SSC14, SSC15, SSC18, and SSCX. The JH-T2T assembly 17 centromeres on chromosomes 1–12, 14, and 16-17 **(Supplementary Table S3**). Putative centromeres were identified in the expected locations in the Sscrofa11.1 chromosome assemblies for SSC1–7, SSC9, SSC13, and SSC18. Two regions harboring centromeric repeats were identified in the chromosome assemblies of each of SSC8, SSC11, and SSC15. Compared to Sscrofa11.1, JH-T2T predicted a greater number of more intact telomeres and centromeres[5].These enhancements highlight the JH-T2T assembly superior quality and utility for genomic research.

By lifting over genes between JH-T2T and Sscrofa11.1, JH-T2T includes 799 newly anchored genes (**Supplementary Table S10**) involved in 96 KEGG entries, enriching two KEGG pathways and 19 GO terms (**Figure S8B** and **Supplementary Table S12**), notably in olfactory (e.g., olfactory transduction) and immunity-related pathways (e.g., cytokine-cytokine receptor interaction, Fc gamma R-mediated phagocytosis, and allergies and autoimmune diseases pathway). Moreover, JH-T2T lost 114 genes (**Supplementary Table 10**), significantly enriching five KEGG pathways and 14 GO terms, including steroid hormone biosynthesis and linoleic acid metabolism (**Supplementary Table S11**).

A comprehensive comparison between the JH-T2T and Sscrofa11.1 has identified 58,200 SVs (28,843 deletions and 29,357 insertions, total genome size of 41.5 Mb, ranged from 50 to 144,010 bp) using Sscrofa11.1 as a reference, with 57,796 medium (50–10,000 bp) and 404 large SVs (≥10 Kb) **(Figure S8C and Supplementary Table S12**). More structural variants were identified in Jinhua pigs than in Ningxiang and Meishan pigs **(Figure S8D**). SVs distribution across chromosomes follow the pattern with higher SVs concentrations in repeat-rich regions (**Figure 3B** and **Supplementary Table S12**). The majority of the SVs (71.23%) were located in repeat regions, suggesting that repeat sequences are an important source of genetic diversity in pigs. They have effectively filled nearly all genome gaps, including the telomeres (**Figure 3A** and **Figure S7**). The majority of these SVs located in intergenic regions (24.08%) and introns (74.65%), with a minority located within coding sequences (CDS) regions (0.24%) (**Figure 3C)**. Moreover, 12,129 genes located in these SVs (**Supplementary Table S12**). Using the pig QTL database, we found SVs enriched in 65 QTLs associated with six economic traits, such as basophil number, drip loss, and head weight (*P-value* <0.01, **Figure S8A** and **Supplementary Table S13**), suggesting SVs potentially impact on important economic traits.

**Figure 3.**
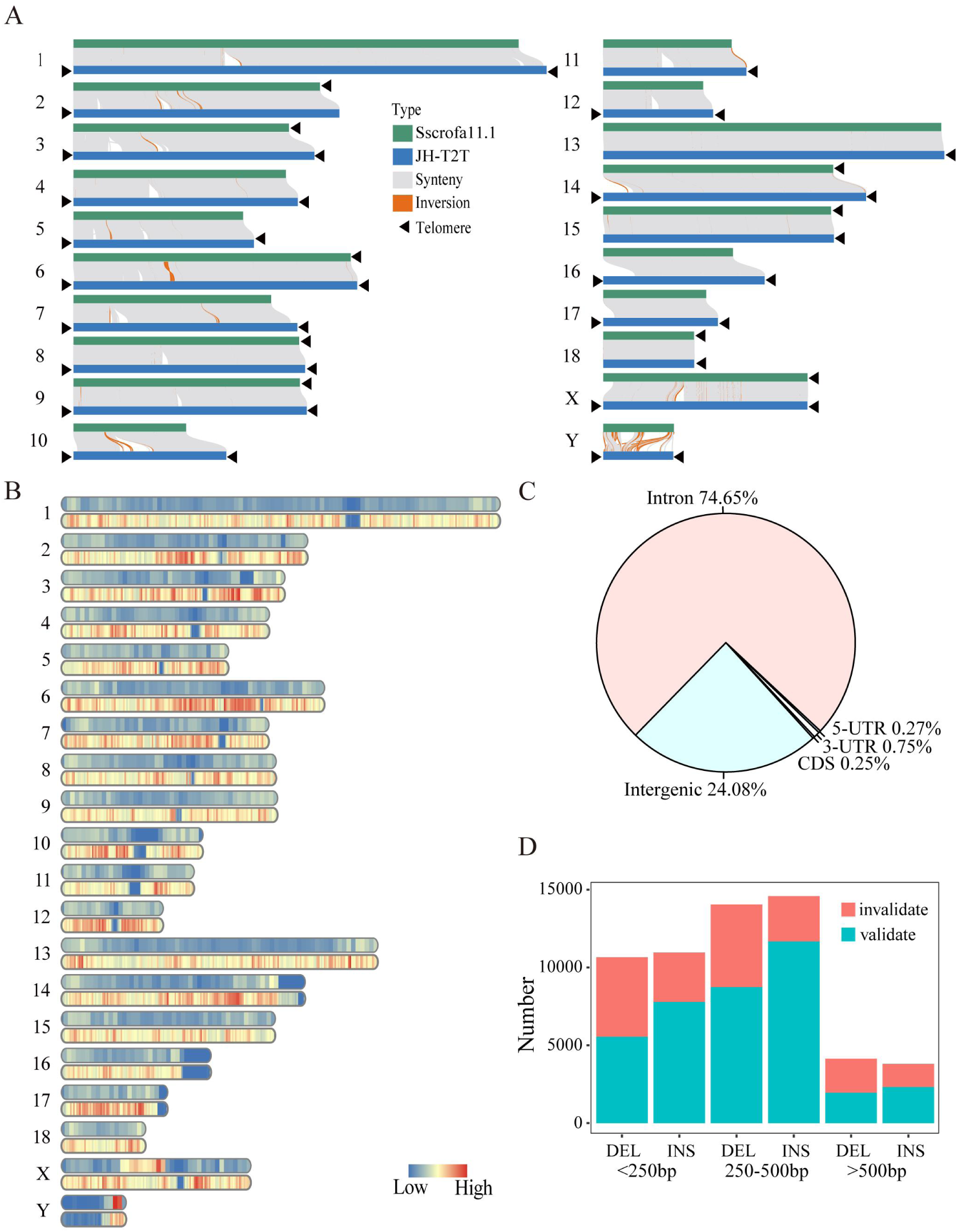
Global comparison of Sscrofa11.1 and JH-T2T genomes. (A) Collinearity between the JH-T2T and Sscrofa11.1 genomes. Collinear regions are shown by gray lines. Black triangles indicate the presence of telomere sequence repeats. (B) Density distribution of SVs across the JH-T2T genome. (C) Proportions of SVs in 5 ‘ UTR, 3 ’ UTR, CDS, introns, and intergenic regions. (D) Percentage of validated SVs categorized by length.

Additionally, we simply validated the detected SVs by examining their sequence coverage using the WGS data from five JH pigs and five Duroc pigs. Employing the JH-T2T and Sscrofa11.1 as refence and applying a validation criterion that required SV mapping sequence coverage greater than 1.00 in one sample and less than 0.90, we confirmed a total of 38,021 SVs (approximal 65.32%), comprising 16,240 DELs and 21,781 INSs, which are associated with 13,967 genes (**Supplementary Tables 14**). Among these, SVs with lengths (>500 bp) were the least frequent (**Figure 3D)**, highlighting the limitations of SV detection through next-generation sequencing data.

### Large-scale genomic differences in JH-T2T genome

In our study, we identified 386 large SVs (≥ 10 Kb) in the JH-T2T genome compared to Sscrofa11.1, including 236 DELs and 150 INSs **(Supplementary Tables S16)**. These SVs affected the presence or absence of 212 genes between the two genomes. Notably, 101 insertions in the Sscrofa11.1 genome contained an additional 100 genes (**Supplemental Table S16**).

Using genotypes from 289 JH pigs and 616 DU pigs (**Supplementary Tables S6**), we analyzed selection signatures to determine if these large SVs experienced selection. By integrating multiple selection detection approaches, such as FST, θπ, and XP-EHH, we identified candidate genomic regions and associated genes overlaps with these SVs. Among these genes, 61 genes located in 96 large SVs showed signals of selection (**Supplementary Tables S17**), being significantly enriched in olfactory transduction (including previously reported pig olfactory transduction genes such as *OR8S1* and novel genes like *OR8B3*, *OR2V2*, and *OR7A17* (**Supplementary Table S18**). The large SVs also harbored genes related to important economic traits. For example, the *CYP2C18* gene, linked to elevated backfat skatole levels in commercial pig populations [56], was located in the largest SV (∼144.0 Kb) on chromosome 14 of JH-T2T, showing signs of positive selection in DU pigs (with low XP-EHH values of JH vs. DU and nucleotide diversity, **Figure 4A-C** and **Supplementary Table S16**). Similarly, an insertion (∼22.2 Kb) in the *GPAM* gene, a marker for intramuscular fat content (IMF) content in musculus longissimus dorsi (MLD)[57] were observed (**Figure 4A and B** and **Supplementary Table S16**). The insertion region showed high FST values and reduced nucleotide diversity, indicating significant genomic differentiation between JH and DU pigs and its potential role in adipogenesis selection in JH pig (**Figure 4D**). The large SVs also contained genes related to immune response, such as *LY9*, *ITLN2*, and *CHIA* (**Supplementary Table S16**). The *LY9* gene region indicated positive selection in DU pigs (**Figure S9B**), associated with immune response regulation [58]. The *ITLN2* and *CHIA* gene reported to link to asthma susceptibility in humans[59].The *ITLN2* gene showed positive selection in DU (**Figure S9C**), while the *CHIA* gene suggested a potential passively positive selection in JH (**Figure S9B**). Those findings may be linked to asthma susceptibility of JH pig. An insertion (∼15.03 Kb) in the *SLA-DOB* gene (**Supplementary Table S15**), which is serve to immune system’s response and relevant to transplant rejection[60].

**Figure 4.**
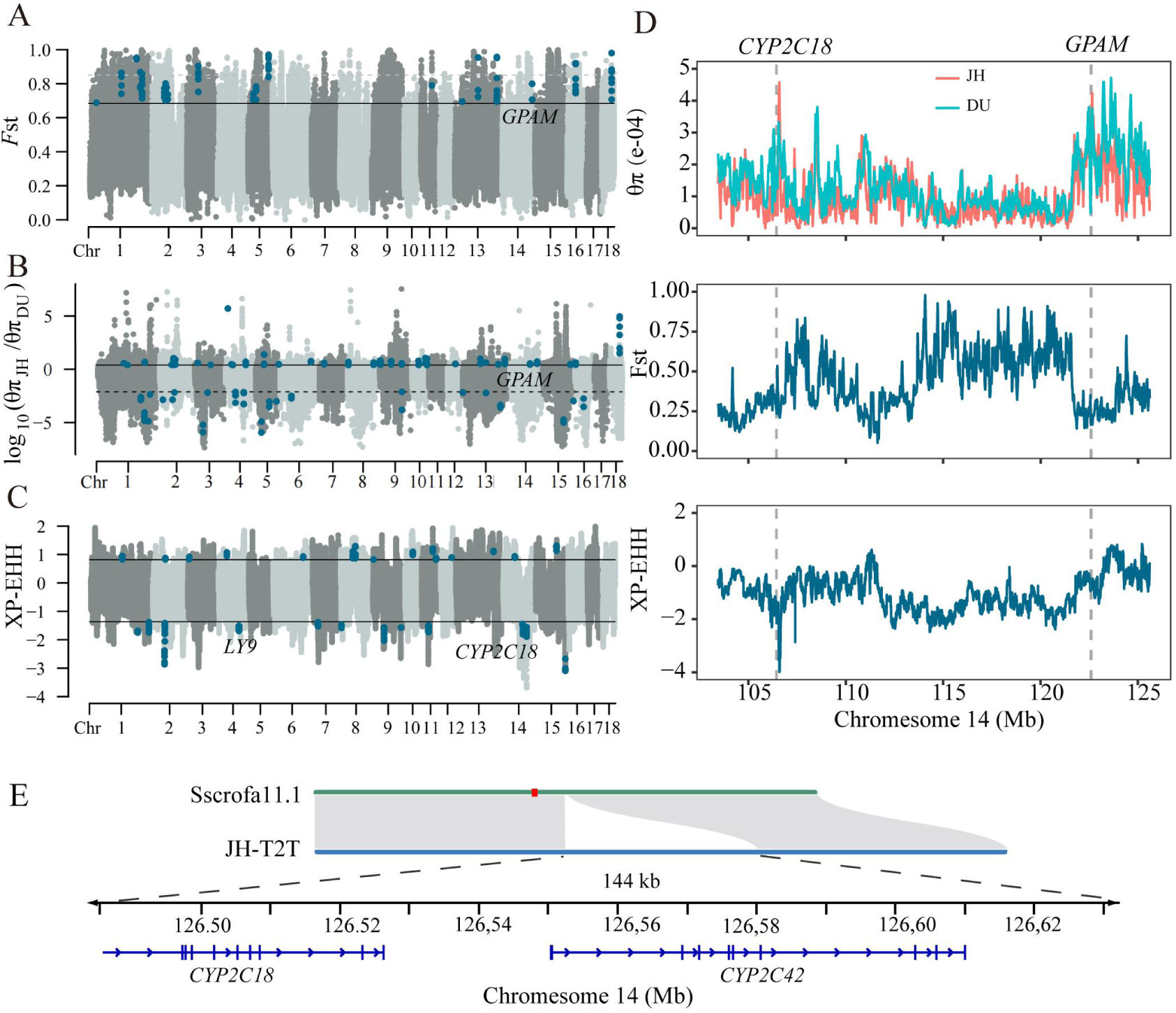
Selection signatures on Large SVs between the Sscrofa11.1 and the JH-T2T genome assembly. (A-C) Genome-wide selection signals between Jinhua and Duroc pigs based on FST, π ratio, and XP-EHH approaches (see detailed in the Supplementary Table S17). (D) FST, π ratio, and XP-EHH in the region of chromosome 14, which contains the genes *GPAM* and *CYP2C18.* (E) The largest SV located in the region of chromosome 14, contained gene *CYP2C42* and *CYP2C18*.

## Discussion

In this study, we built the first T2T pig genome assembly, marking a significant milestone in pig genomics. Our JH-T2T genome assembly demonstrated remarkable improvements over existing assemblies [5–8], both in terms of completeness and quality. Notably, this T2T genome assembly left only six gaps in chromosomes 2, 3, 8, and 10, exceeding the minimum quality standards set by the Vertebrate Genomes Project (VGP) consortium [19].

The high quality of the JH-T2T assembly is evident in its ability to capture complex genomic regions, including repetitive sequences, and telomeres, which were previously inaccessible. This comprehensive coverage addresses the limitations of earlier reference genomes, such as Sscrofa11.1, which contained thousands of gaps and lacked repetitive regions, centromeres, and telomeres. By incorporating these regions, the JH-T2T genome provides a more complete and accurate pig reference genome, essential for detailed genetic studies and breeding programs. Similarly, using the T2T-CHM13 genome yields a more comprehensive view of SVs genome-wide, with a greatly improved balance of insertions and deletions [61].

Advancements in sequencing technology, especially the ONT ultra-long sequencing method, have greatly facilitated the complete assembly of genome. The ONT data played a crucial role in filling gaps, particularly in difficult genomic regions such as repetitive regions, centromeres, and telomeres. Many reference genomes have been successively assembled using ONT reads in farm animals, such as cattle [62], chicken [63] and sheep [64]. In our JH-T2T assembly, ONT reads uniquely filled six out of 63 gaps.

A key advantage of the T2T genome is its superior performance in improving reference genome mapping. The JH-T2T assembly outperforms Sscrofa11.1 in mapping reads from both Western and Chinese pig populations, minimizing gaps and enhancing read alignment accuracy for both DNA and RNA sequencing data. This improvement is crucial for large-scale variant calling from second- and third-generation sequencing data and functional genomics studies, enabling more precise identification of genetic variants and their associated traits. For example, T2T-CHM13 improves the analysis of global genetic diversity based on 3,202 short read-length samples from the 1KGP dataset [61].

Compared to Sscrofa11.1, the JH-T2T genome captures a more comprehensive set of genetic elements. This includes the identification of 799 newly anchored genes not present in Sscrofa11.1, as well as the recognition of 114 genes that were lost in the JH-T2T. This comprehensive capture is made possible by the JH-T2T genome’s ability to fill in gaps and cover repetitive regions, centromeres, and telomeres, which were previously inaccessible. The identification of these novel and lost genes has significant implications for understanding key biological functions, particularly in olfactory function, metabolism, and immune response. Olfactory genes play a critical role in the sensory perception of smell, which is important for behaviors related to feeding, mating, and environmental interaction [65,66]. The JH-T2T genome has enhanced the identification of olfactory receptor genes, such as *OR52B2, OR52B2, OR52B6, OR5K4* and *OR5M3*, providing deeper insights into the genetic basis of olfactory function in pigs. Moreover, genes involved in metabolic pathways are essential for various physiological processes, including growth, reproduction, and overall health. The JH-T2T genome assembly has identified novel genes (such as *ENSSSCG00000026520* and *ENSSSCG00000023320*) related to key metabolic pathways, such as those involved in steroid hormone biosynthesis and linoleic acid metabolism. These genes can influence traits like meat quality and flavor, offering potential targets for genetic improvement in pig breeding. Additionally, the immune system is crucial for protecting pigs against diseases and infections. The JH-T2T genome has uncovered genes (such as *IL1B* and *TLR5* genes) that play important roles in the immune response [67], enabling a better understanding of how pigs respond to pathogens. This knowledge is crucial not only for developing strategies to enhance disease resistance and improve animal health but also for gene editing, producing improved animal models for research, or providing cells, tissues and organ for xenotransplantation.

The comprehensive comparison between the JH-T2T and Sscrofa11.1 has identified 58,200 SVs. Considering that some of the SVs may be due to incomplete genome assembly of Sscrofa11.1, we validated them with WGS data. SVs with lengths (>500 bp) were the least frequent (approximal 65.32%%) of validated SVs may be due to the limited sample size of the WGS data, the validation methodology, or variations in assembly integrity. The JH-T2T genome assembly enables more precise characterization of SVs. This precision is crucial, as incomplete assemblies or technological limitations can result in incorrect assemblies or omissions of important SVs. This may be due to the fact that the sensitivity of second-generation sequencing data for the detection of structural variants is affected by the length of structural variants [68,69]. Since digital PCR (dPCR) has proven to be an excellent tool for sensitive and reliable detection of copy number variation[70], it is hoped that SV differences between JH pigs and Sscrofa11.1 can be further validated in the future by dPCR results.

One of the most critical improvements offered by the T2T assembly is its superior ability to capture SVs that harbor important genes. The JH-T2T genome assembly, with its near-complete coverage, overcomes limitations of earlier reference genomes by providing a comprehensive view of the pig genome especially on poorly characterized regions (e.g., repeat regions) [68,71]. This enhanced coverage allows for the accurate identification and characterization of SVs, which are crucial for understanding genetic variation and its influence on phenotypic traits. For example, among the SVs accurately captured by the T2T genome, we identified notable examples such as the largest SV located in the left telomere region of chromosome 14, which includes important genes like *CYP2C42* and *CYP2C18*. In this study, we systematically characterized large SVs between the two pig genome assemblies, identifying 204 large SVs with gene-model differences. Most of these large SVs overlapped with candidate regions for selection signatures, underscoring the importance of these SVs for pig population differentiation. Some genes likes *LGALS12*, *GPAM*, *CACNB2,*are implicated in essential metabolic pathways and can influence important economically traits [56]. The large SVs also contained genes related to immune response, such as *LY9*, *ITLN2*, and *CHIA*, which may be linked to asthma susceptibility of JH pig [58,59]. The insertion found in the *SLA-DOB* gene, which is serve to immune system’s response and relevant to transplant rejection[72]. When finding ideal organ-source pig with site-specific mutations on SLA to eliminate cross-reacting antibody binding become a strategy [60], it is important to accurately detect variants relevant to immune genes. The ability to accurately detect and analyze such SVs is crucial for advancing our understanding of the genetic basis of these traits. These findings underscored the JH-T2T assembly’s utility in accurately identifying and characterizing SVs, essential for understanding genomic diversity and their potential impact on economic traits and biomedical applications in pigs.

### Conclusions

In conclusion, the JH-T2T genome assembly represents a major leap forward in pig genomics. Its high quality and near-complete coverage significantly enhance our ability to capture and characterize SVs, particularly those harboring important genes. This improvement not only refines the reference genome but also serves as a powerful tool for genetic studies and breeding strategies aimed at improving livestock traits.

## Materials and methods

### Sample collection

The fresh blood was collected from a healthy male Jinhua pig at the National Jinhua Pig Conservation Farm in Zhejiang, China, in 2022 (**Figure 1A** and **Figure S2J**). Ear tissue samples were collected from its parents.

### DNA extraction, library construction, and sequencing

*DNA extraction.* High-molecular weight DNA was extracted using the cetyltrimethylammonium bromide (CTAB) method and purified with the QIAGEN Genomic Kit (Catalog No. 13343, QIAGEN, Hilden, Germany). Ultra-long DNA was extracted using the sodium dodecyl sulfate (SDS) method, omitting the purification step to maintain DNA length. DNA purity was assessed using a NanoDrop One UV-Vis spectrophotometer (Thermo Fisher Scientific). DNA degradation and contamination were monitored on 1% agarose gels. DNA concentration was measured with a Qubit 4.0 fluorometer (Thermo Fisher Scientific).

#### PacBio library preparation and sequencing

SMRTbell target-size libraries were prepared according to PacBio’s standard protocol (Pacific Biosciences, CA) using 15-18 Kb preparation solutions. The main steps included: (1) DNA shearing: high-quality DNA samples (primary band >30 Kb) were selected and randomly fragmented into15-18 Kb pieces using the g-TUBE (Covaris, MA); (2) DNA damage repair, end repair, and A-tailing; (3) Blunt-End ligation: hairpin adapters from SMRTbell Express Template Prep Kit 2.0 (Pacific Biosciences) were ligated; (4) Template purification: imperfect SMRTbell templates were removed with EXOⅢ (from 3’-hydroxyl termini and nicks) and Ⅶ (from 5’-termini) treatment; (5) Size selection: performed using the bluePippin system. Next, the AMPure PB beads were used to concentrate and purify the templates. Then, the sequencing was performed on a PacBio Sequel II instrument with Sequencing Primer V2 and Sequel II Binding Kit 2.0.

#### ONT library preparation and sequencing

Libraries were prepared using the SQK-LSK110 ligation kit following the standard protocol. The purified library was loaded onto primed R9.4 Spot-On Flow Cells and sequenced using a PromethION sequencer (Oxford Nanopore Technologies, Oxford, UK) with 48-h runs at Wuhan Benagen Technology Co., Ltd., Wuhan, China. Base calling of raw data was performed using the Oxford Nanopore GUPPY software (v0.3.0).

#### Hi-C library preparation and sequencing

For Hi-C sequencing, purified DNA was digested with 100 U DpnII and incubated with Biotin-14-dATP. The ligated DNA was sheared into fragments of 300–600 bp, blunt-end repaired, and A-tailed, followed by purification through biotin–streptavidin-mediated pulldown. The Hi-C libraries were quantified and sequenced using the Illumina NovaSeq/MGI-2000 platform.

#### Whole-genome re-sequencing

For whole-genome re-sequencing, total genomic DNA was isolated from fresh blood using the CTAB method. A 150-bp paired-end library with insert sizes of 350 bp was constructed for each individual following standard Illumina library preparation protocols (Illumina). Meanwhile, PCR-free libraries were prepared with the Illumina TruSeq DNA PCR-free library prep kit (Illumina) according to the manufacturer’s instructions. The qualified libraries were then sequenced using an Illumina Hi Seq X Ten platform to produce 150-bp paired-end reads.

### RNA extraction, library construction, and sequencing

For RNA-seq, 19 samples collected form 19 different tissues (hypo, midbrain, hypophysis, surcerebellum, incerebellum, amygdala, pineal, occipital, hippocampus, striatum, parietal, frontal, temporal, muscle, jejunum, ileum, caecum, colon and duodenum) in one JH pig. Total RNA was isolated using the RNAprep Pure Plant Kit (TIANGEN, Beijing, China). All tissues’ total RNA was prepared for mRNA sequencing by using the TRizol reagent. RNA integrity and yield were assessed by the RNA Nano 6000 Assay Kit of the Bioanalyzer 2100 system (Agilent Technologies, Santa Clara, CA, United States) and the NanoPhotometer spectrophotometer (IMPLEN, Westlake Village, CA, United States). For each sample, 3 μg of RNA was used to create sequencing libraries using the NEBNext Ultra TM RNA Library Prep Kit for Illumina (NEB, Ipswich, MA, United States) following the manufacturer’s instructions. Index numbers were added to identify each sample’s sequences. Finally, the clustered libraries were sequenced on an Illumina HiSeq platform, generating 150-bp paired-end reads.

### Datasets and their sources

Genotypes from 939 individuals were collected from PHARP database [21] **(Supplementary Table S6).** Additionally, 92 RNA-seq data from ten pig population, covering eleven different tissues (brain, heart, liver, spleen, lungs, kidneys, fat, muscle, ovaries, testicles, and intestinal segments) were downloaded from NCBI **(Supplementary Table S7)**.

### Genome size estimation

To estimate the pig genome size and address potential issues such as sister chromatid merging and repetitive sequences, we used k-mer analysis with the jellyfish software (version 2.2.10) [22]. The command “Jellyfish count -G 2 -m 17 -C” and “histo kmercount” were used to calculate the k-mer count and generate histograms, respectively.

### Genome assembly

The main goal of this study was to create a high-quality, gapless assembly of the Jinhua pig, comprising 18 autosomes and two sex chromosomes (X and Y). The assembly process followed the Vertebrate Genomes Project (VGP) assembly pipeline [23] with modifications (**Figure 1A** and **Figure S1**). First, the initial assembly was constructed using PacBio HiFi reads and ONT ultra-long reads (**Supplementary Table S1**). For the PacBio assemblies, consensus reads (HiFi reads) were generated using CCS software (https://github.com/PacificBiosciences/ccs) with the default parameter. HiFi reads were then assembled using Hifiasm (version 0.16.1-r375) [15,24]. ONT reads were assembled using NextDenovo (version 2.5.0) [25] (**Supplementary Table S1**). Second, an auxiliary assembly was performed using Allhic [26] and juicebox [27] to improve the Hifiasm output assembly with the help of Hi-C reads. Allhic was utilized to assign the assembled contigs/scaffolds to near-chromosome level. The chromosome interaction intensity, based on the juicebox software, was used for manual correction.

### Gap filling

To fill the gaps in the genome assembly, we used the winnowmap (v1.11) software with parameters (k=15, -MD) [25]. This process involved comparing the hole-filling data (error-corrected ONT genome versions, HiFi or ONT reads) with the genomic gap intervals. The priority for gap filling steps was given first to error-corrected genome versions, followed by ONT and HiFi reads. Using this approach, we reduced the number of gaps from 63 to 14. The remaining 14 gap regions could not be adequately covered by the assembly/ONT/HiFi data due to a lack of good reads. We then mapped these gap regions with Hi-C data, generated Hi-C interactions, and imported them into juicebox. After identifying mapping errors, we made manual adjustments, resulting in a final JH-T2T genome with only six gaps **(Figure S3)**.

### Genome assembly quality assessment

To systematically evaluate the quality of the genome assembly, we conducted the following assessment: i) Gene completion. The gene completion of the assembly was evaluated using BUSCO (v5.4.3) with the mammalia_odb10 dataset [25]. ii) Genome continuity. The genome continuity was assessed by calculating contig N50 length using QUAST (v5.0.2) [29]. iii) Quality value (QV). Merqury [30] was used to calculate QV combining Illumina reads. iv) Reads mapping rate and coverage. We mapped the WGS (n=153), and HiFi (n=1) and ONT (n=1) reads to the assembly using BWA-MEM2 and minimap2 [31], respectively. We then calculated their mapping rates and coverages.

### Identification of telomeres and centromeres

In vertebrates, telomeres consist of conserved repetitive sequences as described in the Telomere Database (http://telomerase.asu.edu/sequences_telomere.html). Here, we also used the vertebrate telomeric repeat (6-mer TTAGGG/CCCTAA) to identify telomeres using the Tidk (v0.2.0) tool [32]. Centromics software (https://github.com/ShuaiNIEgithub/Centromics) was used to pinpoint centromere regions. This tool utilizes characteristics such as a high density of short tandem repeats and a low density of genes, which are typical of centromere regions, to identify centromeres in the JH-T2T genome.

### Repeat annotation

The homologous repeat annotation library for the JH-T2T genome was constructed by extracting mammalian repeat sequences from a combined library comprising Repbase (release 20181026) and Dfam (version 3.2)[33,34]. RepeatModeler (version 2.0.3)[35] was then used to analyze and predict repeat sequences based on this library. Finally, the Repeatmasker (version 4.1.2)[36] was employed to annotate the transposable elements (TEs) in the JH-T2T genome using the custom non-redundant set of repeats.

### Gene annotation

To annotate the protein-coding genes in the JH-T2T genome, a combination of ab initio, homology-based, and transcriptome-based prediction methods were employed. For the ab initio gene prediction, the MAKER3 pipeline [37] was applied to predict gene structures in the masked JH-T2T genome. High-quality protein sequences from Ensembl release 106 were used for gene annotation, including homologous protein sequences from eleven closely related species (*Homo sapiens, Equus caballus, Canis lupus, Bos_taurus, Capra_hircus, Ovis_aries, Camelus dromedaries, Delphinapterus leucas, Balaenoptera musculus, Physeter catodon, and Tursiops truncates*). Additionally, transcripts from 111 samples (**Supplemental Table S7**) generated from our RNA-Seq data and public available data were processed using HISAT2 (v2.2.1) and StringTie (v2.1.4) [38,39]. The initial round of gene annotation utilized protein sequences and transcripts. BLASTN [40] with an e-value cutoff of 1e-10 was used to map these homologous protein sequences to the JH-T2T genome. Only the protein sequences with the highest-scoring alignments, having a minimum identity score greater than 80%, were retained to predict putative gene models using Exonerate (v2.4.0) [41]. The second round transcript-based gene prediction involved training SNAP (v2006-07-28) [42] and AUGUSTUS (v3.4.0) [43] with predicted gene models to predict genes.

### Functional annotation of protein-coding genes

We employed three methods to annotate functions of protein-coding genes. First, protein sequences similarity were searched against the NCBI nonredundant protein database and the Swiss-Prot database [44,45] using BlastP software [40]. Second, protein domain and gene ontology term annotations were performed using InterProScan [46]. Third, KEGG annotation was performed with the kofam_scan [47]. These methods provide complementary approaches, combining sequence similarity, domain analysis, and pathway information to gain insights into the potential functions of these genes in the JH-T2T genome. Additionally, the expression of these genes were also examined using the RNA-seq data. We first used the fastp [48] to remove the low-quality reads and adapters in the raw RNA-seq reads, and mapped the remaining reads to the transcripts of high-quality predicted genes by Hisat2 [38]. We then used StringTie (v.2.1.7) [39] to assemble and quantify transcripts guided by the JH-T2T genome. The transcripts were evaluated based on transcripts per million (TPM) values. A TPM > 0 indicated the presence of a transcript in a sample. If a transcript occurred in at least one sample, it was considered as validated, indicating the expression of the predicted gene.

### Global comparison of the Sscrofa11.1 and JH-T2T genome

To assess variation in chromosome-scale synteny, we compared the JH-T2T and Sscrofa11.1[5] assemblies. We began by aligning the two genomes using NUCmer [49] with parameters −l 100 -c 1000, refining the results with Delta-filter using parameters -i 95 −l 100 −1. Additionally, Minimap2 [31] with parameters -cx asm5 -t8 --cs was used to align Sscrofa11.1 to JH-T2T. The optimal alignments were used for SNPs and indels calling with paftools.js [31]. For detecting structural variants (SVs), we used Minimap2 with parameters -a -x asm5 --cs -r2k to to get the best alignments, followed by SV calling with svim-asm using the haploid parameter [50]. To explore the functional implications of deleterious variants, we selected genes with such variants for enrichment analysis using KOBAS [51]. Next, we employed Liftoff (version 1.6.2) [52] to map genes between Sscrofa11.1 and JH-T2T, assessing their consistency. We also aligned WGS clean reads including JH and Duroc pigs to both assemblies using BWA-MEM tool with default parameters [53] to examine the coverage and depth of detected SVs. These analyses allowed us to assess variation in chromosome-scale synteny, identify genetic variants, investigate missing genes, and validate SVs in the JH-T2T and Sscrofa11.1.

### Selection signatures between JH and Duroc pigs in large SV regions

To examine selection signatures in large SV regions between Jinhua and Duroc pigs, we analyzed genotypes from 289 Jinhua and 616 Duroc pigs **(Supplemental Table S6)**. We used three approaches to detect selection signals: fixation index (FST), nucleotide diversity ratio (θπ), and cross-population extended haplotype homozygosity (XP-EHH). The FST and θπ were calculated across the genome using 10 Kb non-overlapping sliding windows with VCFtools (v0.1.16) [54]. The XP-EHH was conducted with selscan (v1.2.0) [55], averaging XP-EHH scores over 10 Kb non-overlapping sliding windows. Genomic regions in the top 5% values for at least one selection signature were identified as selective sweeps. Genes in these selective sweep regions were considered candidate high-related genes.

### Ethics approval and consent to participate

Ethical permission to collect blood samples from pigs was approved by the Institutional Animal Care and Use Committee of Zhejiang University. All procedures in which pigs were involved were per the agreement of the Institutional Animal Care and Use Committee of Zhejiang University (ZJU20220262).

## Consent for publication

Not applicable.

## Availability of data and materials

The genome assembly for the Jinhua pig (JH-T2T) is available at http://alphaindex.zju.edu.cn/ALPHADB/download.html. The genotype datasets generated and/or analyzed during the current study are available at PHARP (http://alphaindex.zju.edu.cn/PHARP/index.php) and at the SRA repository (https://www.ncbi.nlm.nih.gov/sra). See the ‘ MATERIALS AND METHODS ’ section above for their availability. Computer code for data processing is available from the authors upon request.

## Authors’ contributions

Z.W. and YC.P. conceived and supervised the study. CY.C. and Z.W. wrote the manuscript. CY.C. performed the majority of the analyses. J.M., JB.S., QQ.X., H.C., ZY.Z., F.W., S.L. and XW.Y. prepared the DNA sampling and experiments. Z.Z. and QS.W. participated in the discussion of the results. All authors read and approved the final manuscript.

## Acknowledgments

We thank all the researchers worldwide that made their sequencing data publicly available.

## Supplementary Tables

Supplementary Table 1. Summary of the sequence data used for JH-T2T assembling.

Supplementary Table 1. Statistics on the number and length of clusters of individual chromosomes and Genomic mount rate.

Supplementary Table 3. Genome statistics, predicted telomeres and centromeres.

Supplementary Table 4. Scaffold and contig length of four assemblies.

Supplementary Table 5. BUSCOs analysis of JH-T2T, Sscrofa11.1, MS, NX.

Supplementary Table 6. Information of the 939 WGS pigs.

Supplementary Table 7. Information of the 111 RNA pigs.

Supplementary Table 8. Gap positions of JH-T2T.

Supplementary Table 9. Summary of repeat content of JH-T2T.

Supplementary Table 10. Lost and gain genes between JH-T2T and Sscrofa11.1.

Supplementary Table 11. Newly gain genes in KEGG PATHWAY and Gene Ontology.

Supplementary Table 12. List of SVs between JH-T2T and Sscrofa11.1.

Supplementary Table 13. Statistics of SV-related QTL.

Supplementary Table 14. SV with WGS validation.

Supplementary Table 15. Large SVs localized on Sscrofa11.1 reference and overlapping gene ID on Large SVs.

Supplementary Table 16. Selected large SVs localized on Sscrofa11.1 reference and overlapping gene ID on Large SVs.

Supplementary Table 17. Novel predicted genes in KEGG PATHWAY and Gene Ontology.

## Supplementary Figures

Figure S1. Overview of the data processing pipeline used for the assembly and genomic analysis of JH-T2T genome.

Figure S2. Summary of data used for JH-T2T assembly.

Figure S3. Misassembly correction using Hi-C data.

Figure S4. Evaluation of the JH-T2T assembly.

Figure S5. Replaced gap regions.

Figure S6. Coverage of WGS, HiFi, and ONT read on 47 filled gaps region of the JH-T2T assembly.

Figure S7. Predicted centromeres’ locations in the JH-T2T assembly.

Figure S8. SVs between the Sscrofa11.1 and JH-T2T.

Figure S9. Selective regions on Large SVs between the Sscrofa11.1 and the JH-T2T genome assembly.

