## Supplementary Materials for "The first near-complete genome assembly of pig: enabling more accurate genetic research"

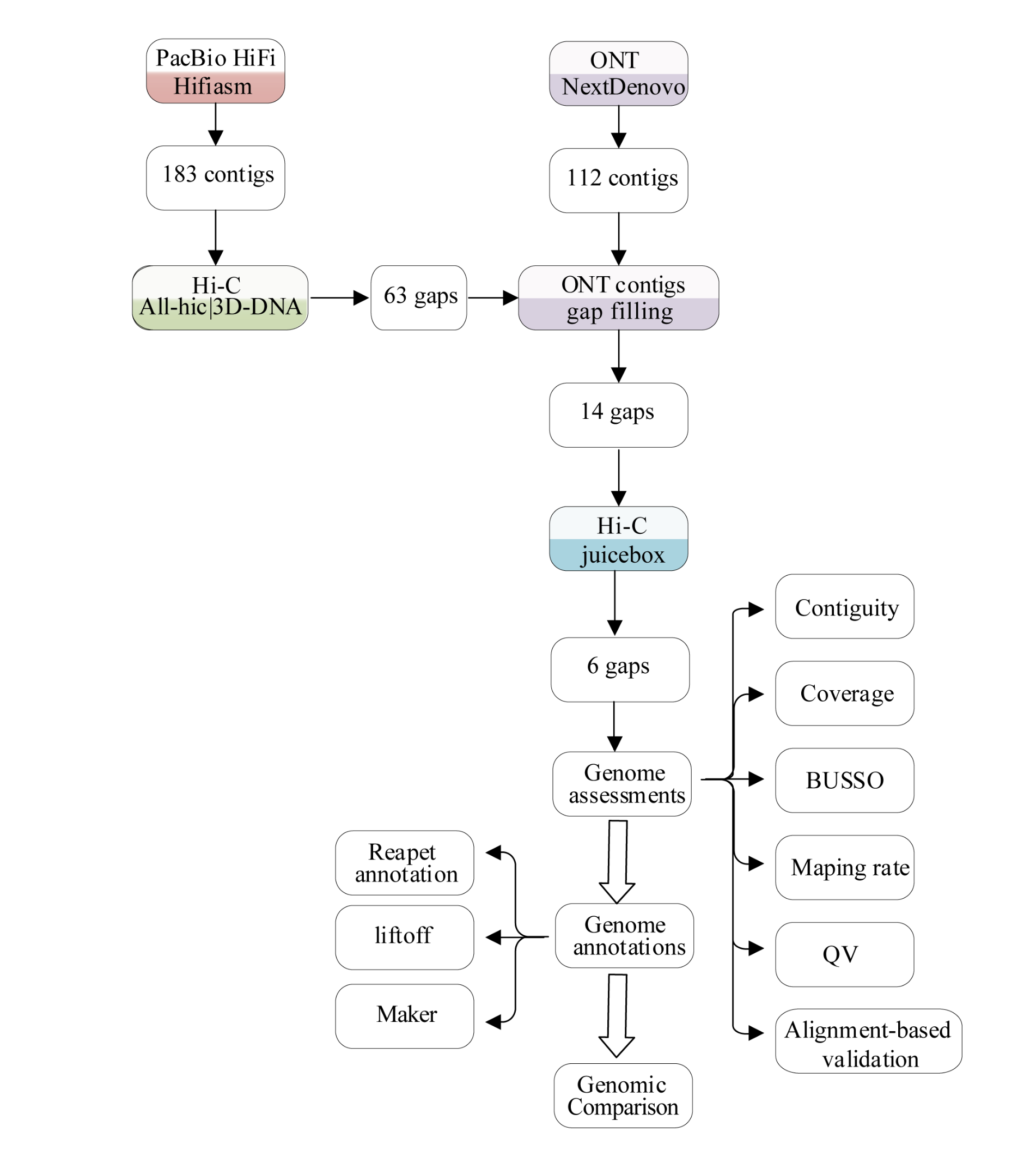

**Figure S1. Overview of the data processing pipeline used for the assembly and genomic analysis of JH-T2T genome.**

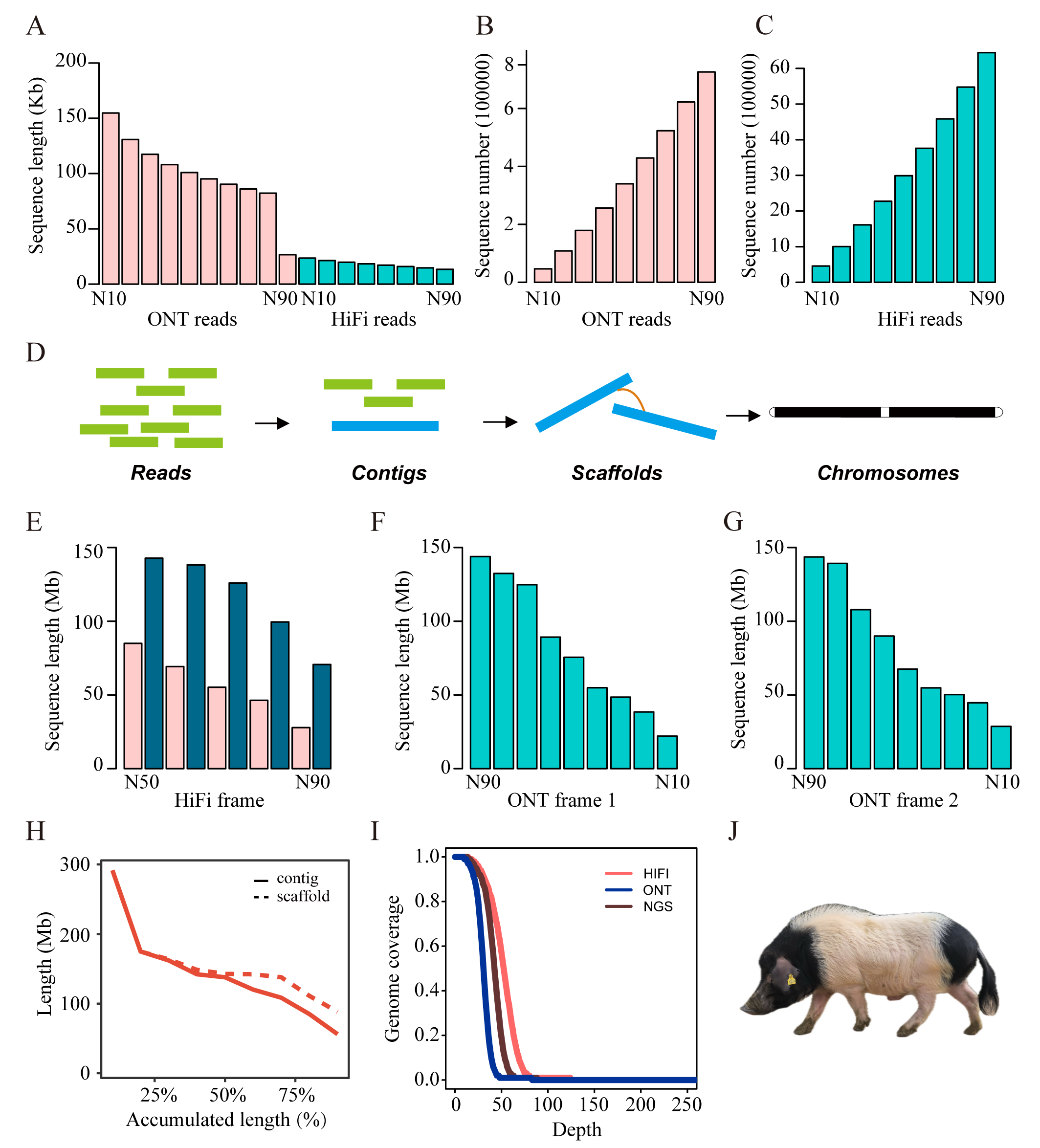

**Figure S2. Summary of data used for JH-T2T assembly.** (A) Length distributions of ONT ultra-long and PacBio HiFi reads. Each bar indicates the length rang from read N10-N90. (B-C) Read count distributions of ONT ultra-long and PacBio HiFi reads. Each bar indicates the count of read from N10 to N90 per100000. (D) Genome assembly pipeline. Overview of the steps and processes involved in assembling the JH-T2T genome. (E) Length distributions of PacBio HiFi contigs before and after Hi-C interaction. Pink bars depict contig lengths before Hi-C interaction, while blue bars show lengths after Hi-C interaction. (F) Length distributions of ONT contigs using only ONT ultra-long reads with a read N50 of 100 Kb. (G) Length distributions of ONT contigs using all ONT ultra-long reads (read N50 52.17 Kb). (H) Continuity of ultra-long reads assembled contigs. Assessment of the continuity and completeness of contigs assembled from ultra-long reads. (I) Read coverage of JH-T2T by ONT ultra-long reads, HiFi reads and NGS reads. (J) A healthy male Jinhua pig at the National Jinhua Pig Conservation Farm in Zhejiang, China.

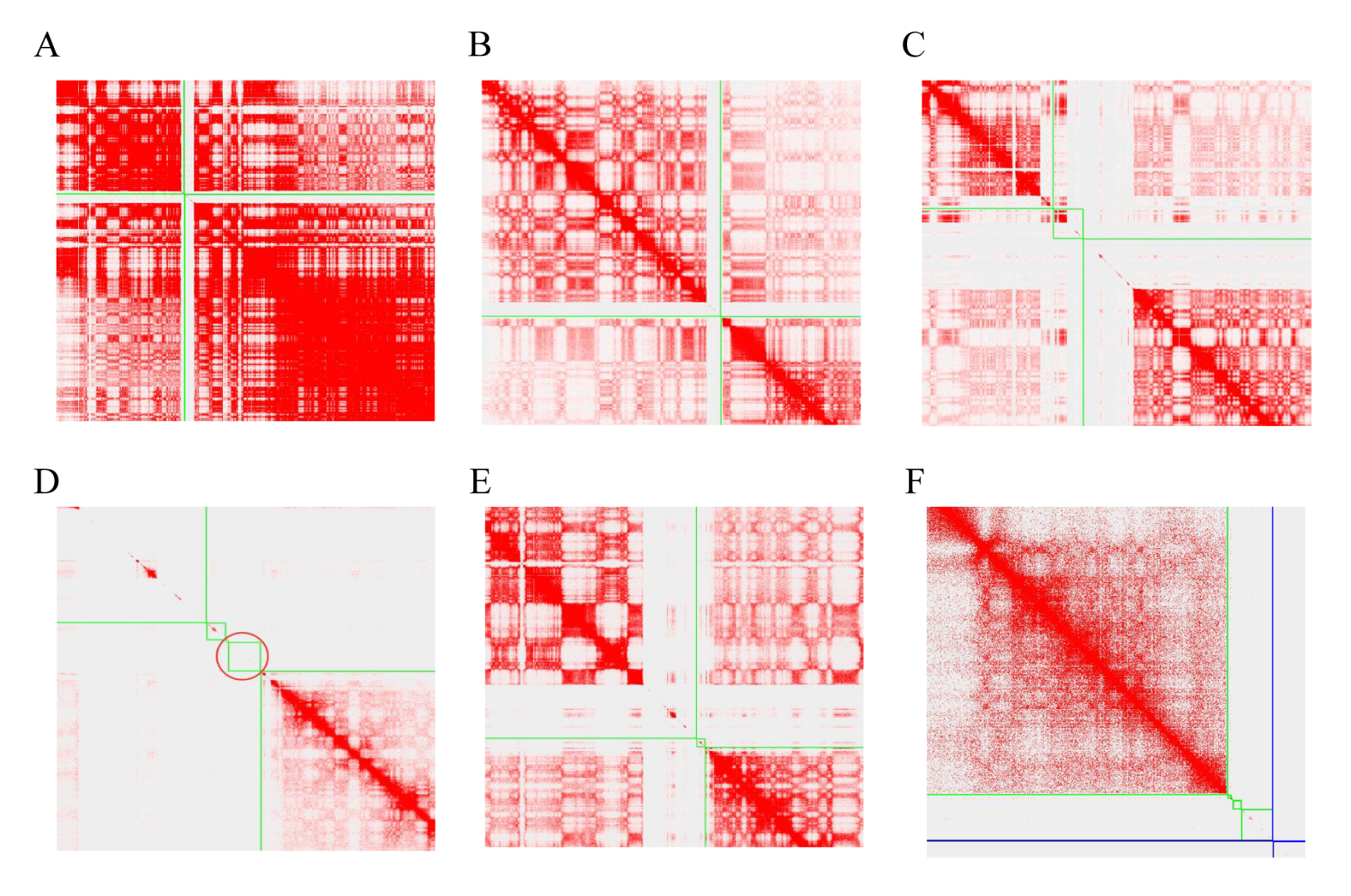

**Figure S3. Misassembly correction using Hi-C data.** (A-C) Hi-C map of Chromosomes 2, 8, and 13 before correction. Illustrates gaps in Chromosomes 2, 8, and 13 of the JH-T2T assembly that was identified through scaffolding without subsequent editing. (D) Hi-C map of Chromosome 3 after correction. Displays Chromosome 3 following the correction of misjoins, with the previously erroneous joins highlighted by red ellipses. (E) Hi-C map of Chromosome 13 after correction. Presents Chromosome13 after the misjoins have been rectified, with corrected areas indicated by red ellipses.

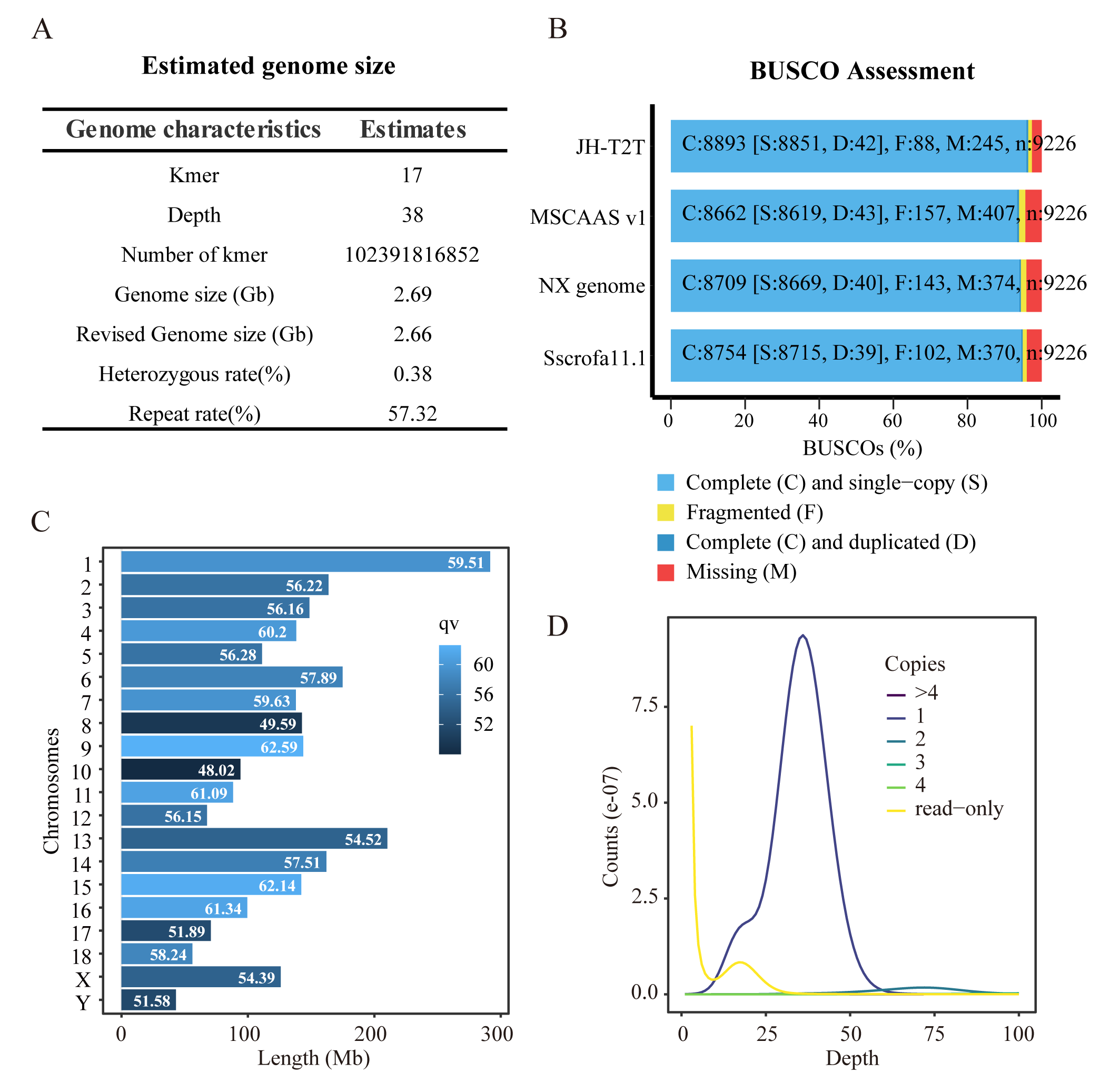
**Figure S4. Evaluation of the JH-T2T assembly.** (A) Estimated genome size of Jinhua pig. (B) BUSCO assessment of JH-T2T, Ninxiang, Meishan, and Sscrofa11.1 genomes. (C) QV distribution across Chromosomes of JH-T2T. Bar plot displaying the Quality Value (QV) metrics for each chromosome within the JH-T2T assembly. QV scores indicate the accuracy of the base calls, with higher values representing fewer errors. (D) Copy number spectrum (spectra-cn) of k-mers for JH-T2T. The plot focuses on k-mers with copy numbers greater than one (i.e., 2, 3, 4, and >4), which are presented in an enlarged view to emphasize repeat regions and structural variations within the genome..

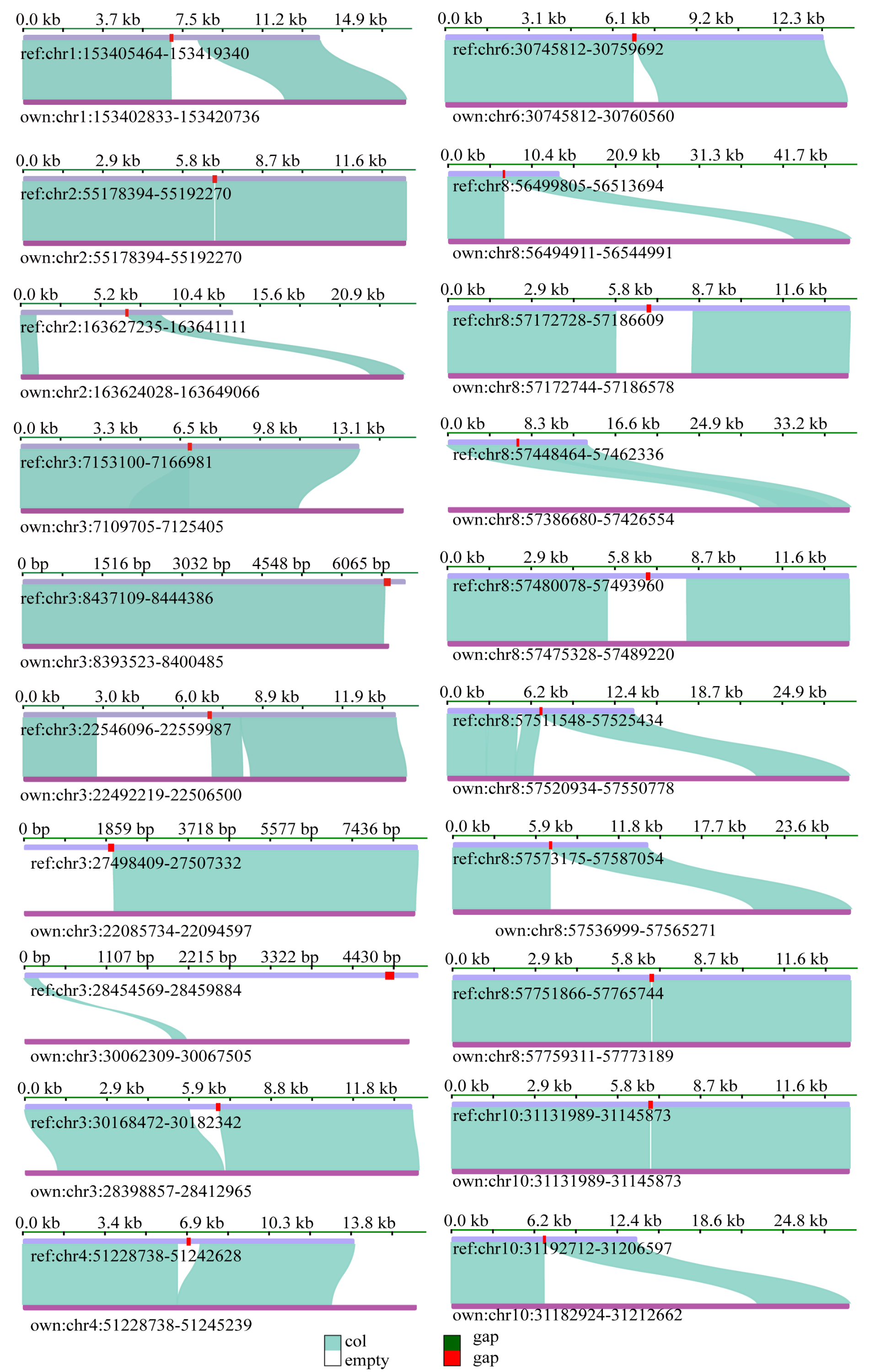

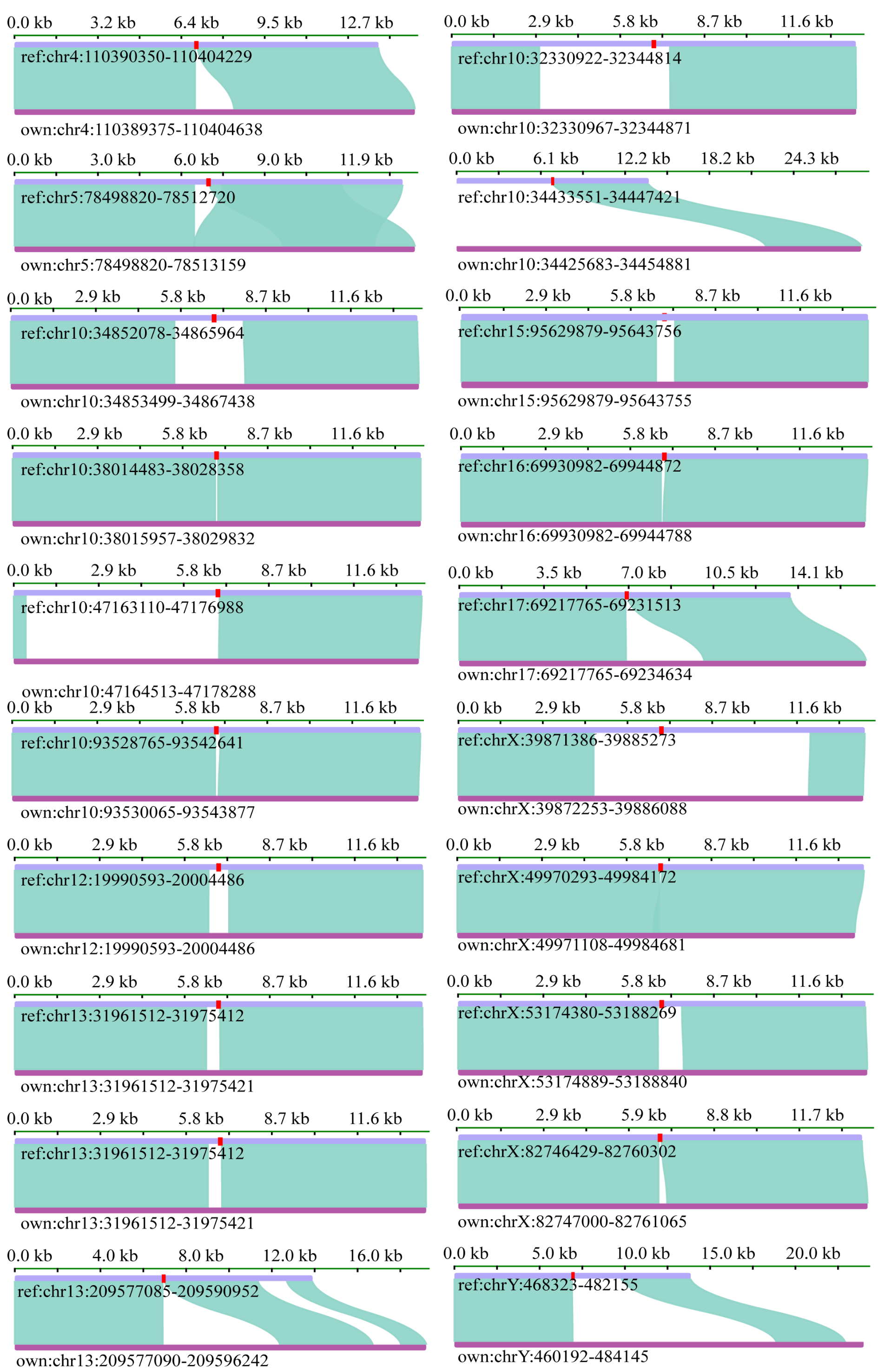

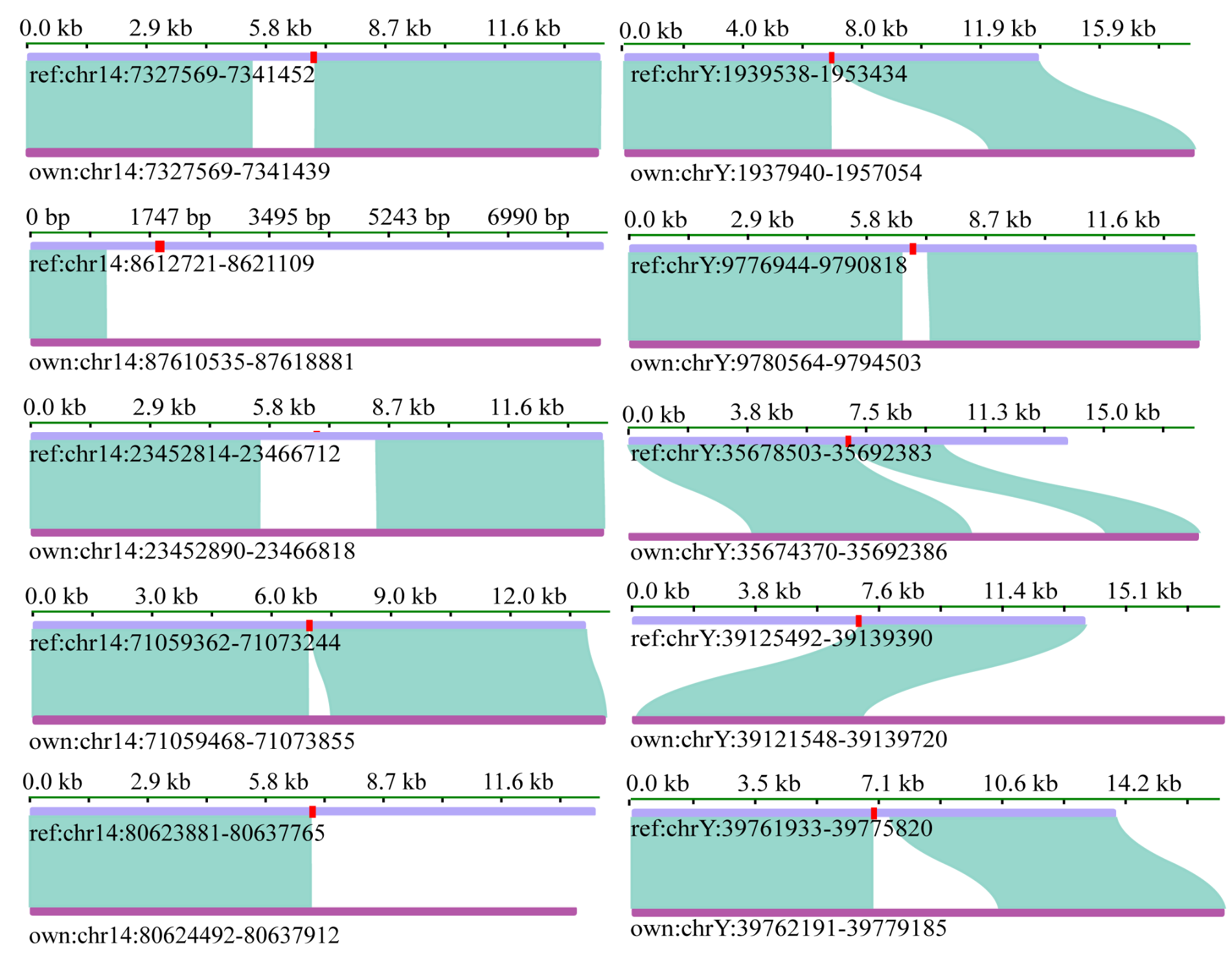

Figure S5. Replaced gap regions.

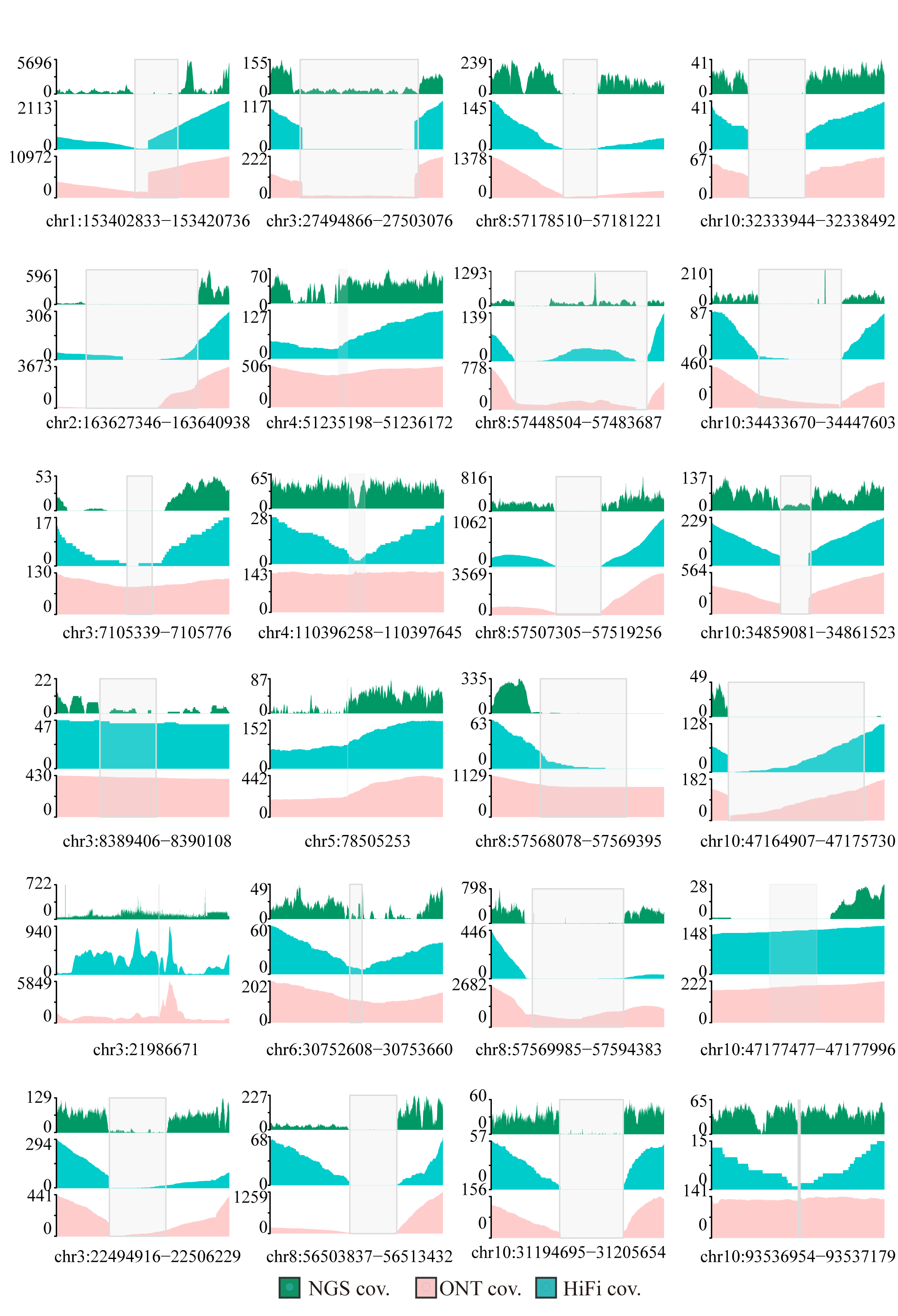

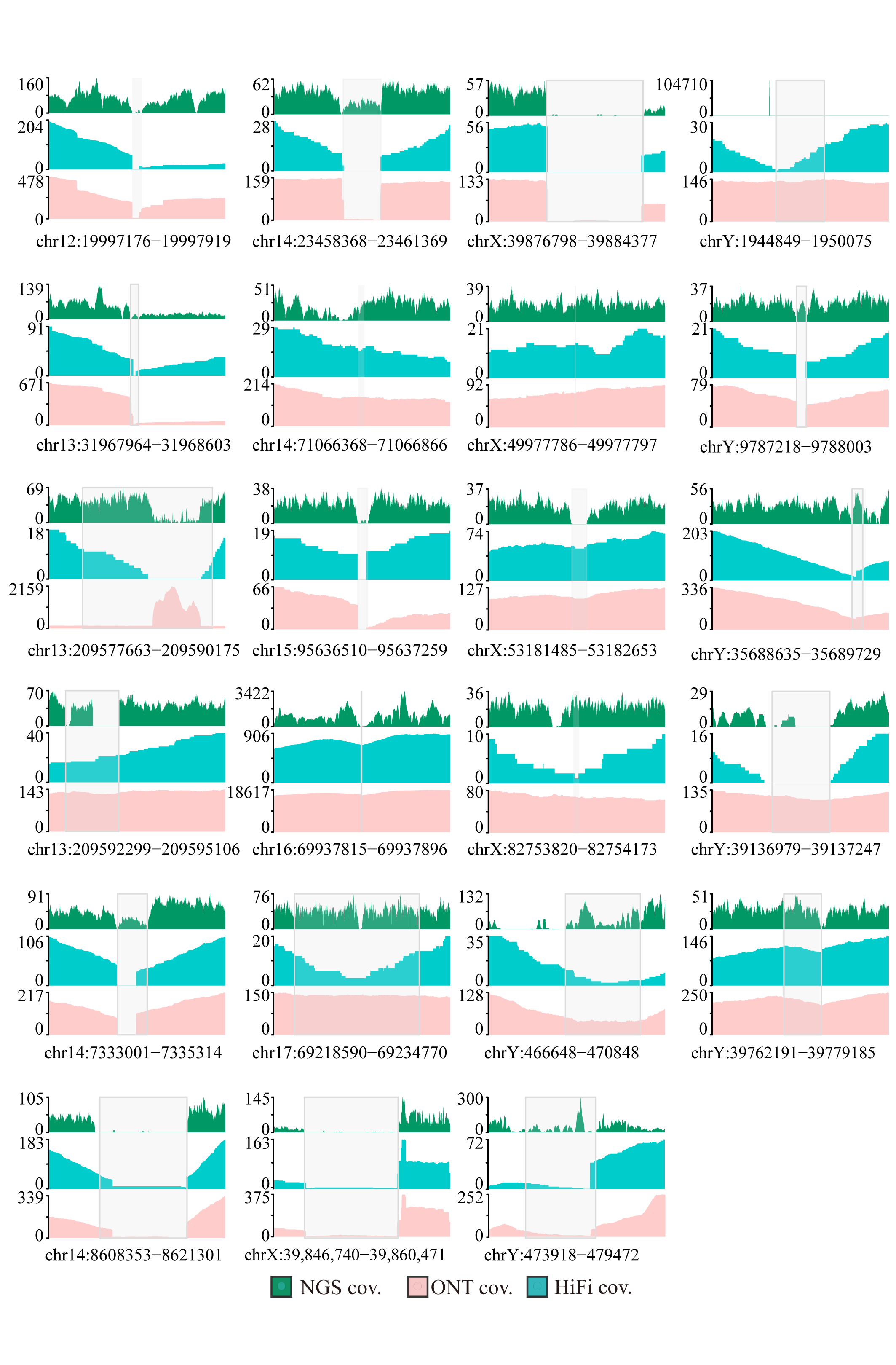

Figure S6. Coverage of WGS, HiFi, and ONT reads on 47 filled gaps regions of the JH-T2T assembly. The x-axis represents chromosome position, and the y-axis represents read sequencing depth.

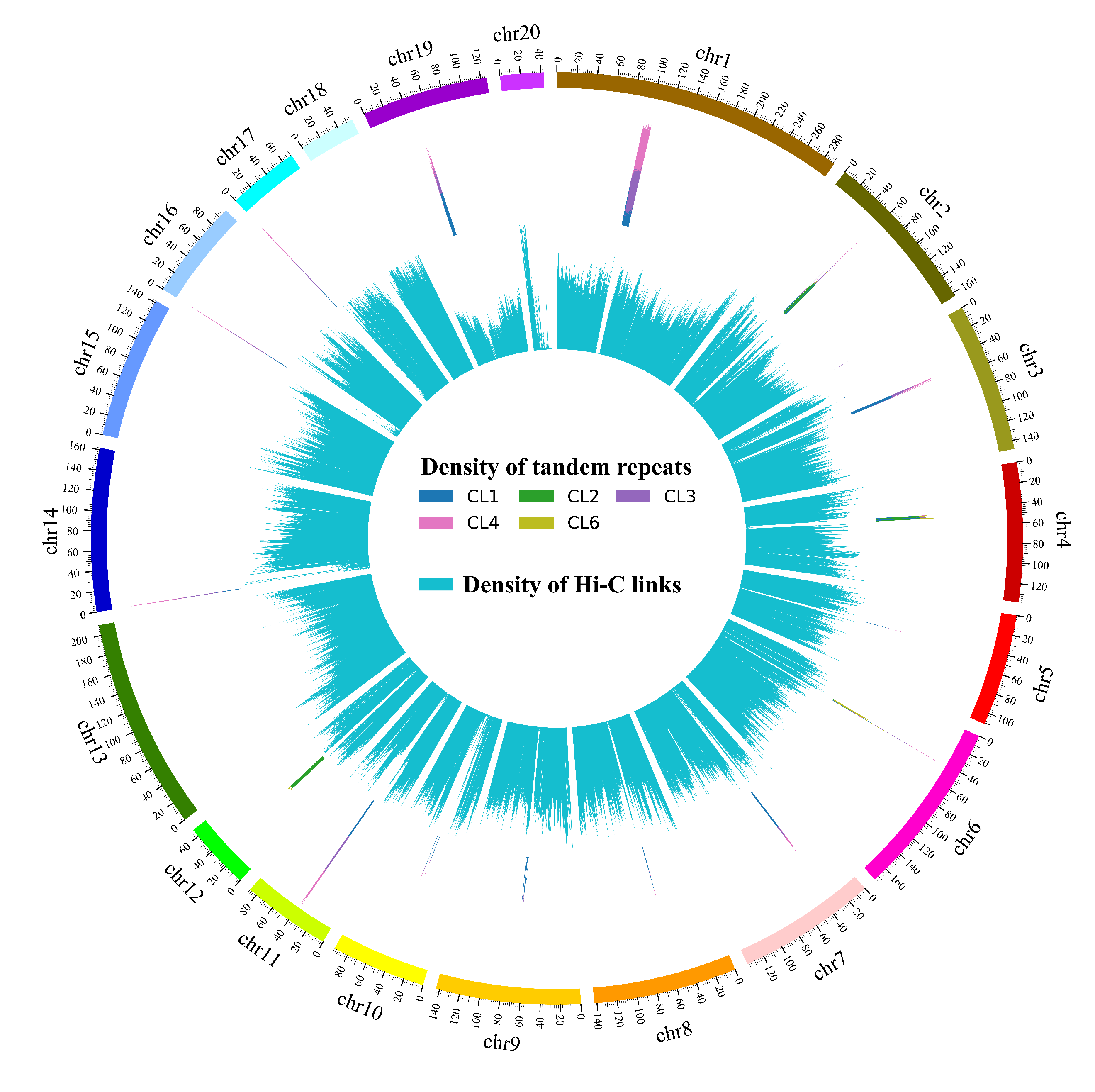

Figure S7. Predicted centromeres’ locations in the JH-T2T assembly.

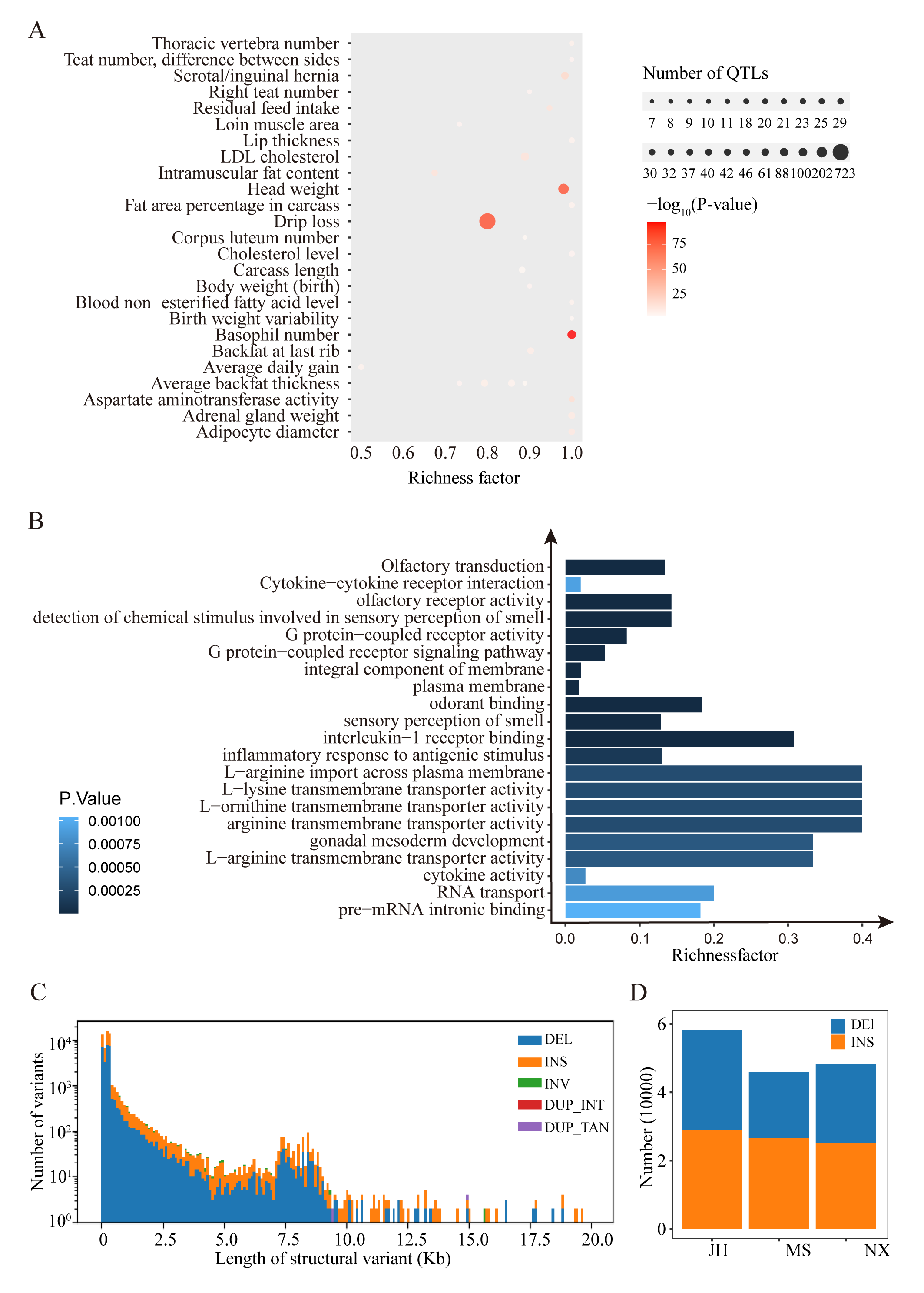
Figure S8. SVs between JH-T2T and Sscrofa11.1. (A) Proportions of different QTL classes where SVs occurred. (B) Novel predicted genes in KEGG PATHWAY and Gene Ontology. (C) Length distribution of SVs. (D) Numbers of SVs in JH-T2T, MS, MX compared to Sscrofa11.1.

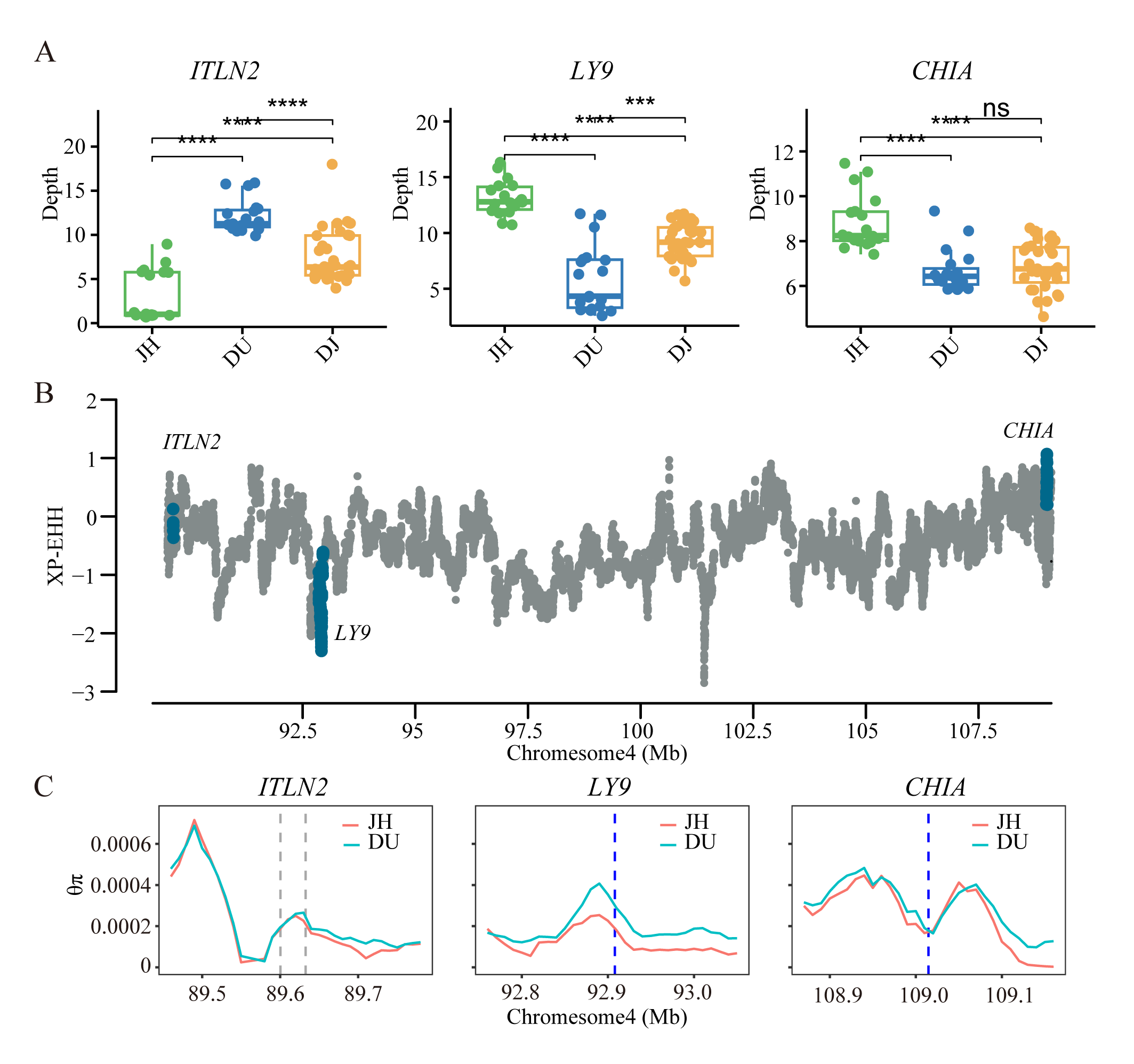

Figure S9. Selective regions on Large SVs between the Sscrofa11.1 and JH-T2T genome assemblies. (A) The sequencing read depth across specific regions within large structural variants (related gene *ITLN2*, *LY9* and C*HIA*)*.* (B) Selective region (related gene *ITLN2*, *LY9* and C*HIA*) between Jinhua and Duroc pigs based on XP-EHH approaches. (C) Selective region (related gene *ITLN2*, *LY9* and C*HIA*) between Jinhua and Duroc pigs based on π ratio approaches.
